# FACT safeguards promoter topology by maintaining nucleosomes and restricting chromatin factor spreading

**DOI:** 10.64898/2026.02.18.706382

**Authors:** Ana M. Dopico-Fernandez, Hangpeng Li, Catherine Chahrour, James L.T. Dalgleish, James O.J. Davies, Robert A. Beagrie, Thomas A. Milne

## Abstract

Facilitates chromatin transcription (FACT) is a histone chaperone that displaces and re-assembles histones during transcription. Recent studies have reported a minor role for FACT in chromatin architecture. We have recently shown that active gene promoters form nanoscale domains and proposed they are created by the biophysical properties of nucleosome-free regions. Here we use base-pair resolution Micro Capture-C ultra to show that, following FACT degradation, nanoscale domains are lost and subnucleosomal chromatin interactions are rearranged at active promoters. Nucleosome-free regions at these promoters expand and chromatin-binding factors invade the newly accessible chromatin, indicating FACT maintains the integrity of active promoters by opposing DNA-binding factor spreading into gene bodies. Finally, we show increased interactions between promoters across topologically associating domains, suggesting large-scale structural changes upon FACT loss. Thus, we demonstrate FACT plays a major role in chromatin organisation and provide *in vivo* evidence that nucleosomes drive both local and long-range chromatin architecture.

## INTRODUCTION

The eukaryotic genome exists in the form of chromatin packed into different levels of organisation. Advances in high-resolution microscopy and chromosome conformation capture (3C) techniques have demonstrated that this three-dimensional organisation is tightly controlled, and its proper regulation is crucial for DNA-templated processes, including the temporal and spatial regulation of gene expression.

The fundamental subunit of chromatin is the nucleosome—an octamer of histone proteins with 147 bp of DNA wrapped around it. Nucleosomes are connected by linker DNA into nucleosome arrays, which further interact with one another forming chromatin fibers. At the sub-megabase scale, chromatin folds into topologically associating domains (TADs)^1^ that contain interacting regulatory elements (e.g. promoters and enhancers). At many sites, TAD formation is thought to be mediated by loop extrusion, in which chromatin progressively threads through the ring-like cohesin complex until it is stalled at boundary elements^2–4^. At a larger scale, chromatin is segregated into A and B compartments, which are respectively enriched in euchromatin and heterochromatin^5^.

Chromosome conformation capture approaches that capture all pairwise contacts within an area (e.g. Hi-C^5^, Micro-C^6–8^, 5C^9^, tiled MCC^10^ and Region Capture Micro-C^11^) have been instrumental in identifying these principles. However, genome-wide techniques become impractically expensive at higher resolution, and even the highest-resolution datasets miss key information about nucleosome organisation and bound protein complexes as they cannot resolve subnucleosomal features. Our understanding of the mechanisms driving chromatin structure and distal regulatory interactions has therefore been limited by the inability to resolve chromatin structure at base-pair resolution.

To overcome this limitation, we recently developed Micro Capture-C ultra (MCCu), a tiled 3C technique that allows us to resolve chromatin structure and cis-regulatory interactions at base-pair resolution^12–14^. The use of tiled capture probes captures contacts from all interacting fragments within the region of interest, which can be used to construct multidimensional contact maps and visualise 3D chromatin structure at subnucleosomal resolution. Our initial MCCu results, combined with super-resolution microscopy and molecular dynamics simulations, led us to propose a model in which genome structure is driven by the biophysical properties of chromatin^14^.

In this model, chromatin at the kilobase level is partitioned by nucleosome-free regions (NFRs) into globule-like structures termed nanoscale domains. Molecular dynamics simulations indicate that nanoscale domains are largely driven by the intrinsic biophysical properties of nucleosomes: unacetylated nucleosomes form a core, acetylated nucleosomes are enriched in the periphery and NFRs separate from aggregated nucleosomes.

Our model integrates new super-resolution microscopy data with older electron microscopy and 3D super-resolution imaging^15–17^, which found that the nucleus is divided into the “inactive nuclear compartment” (containing densely compacted heterochromatin) and the “interchromatin space” (rich in diffusible proteins and RNA) separated by the “perichromatin region” (containing loosely compacted euchromatin). In our model, the inactive nuclear compartment corresponds to the gel-like core of unacetylated nucleosomes, the perichromatin region to the peripheral acetylated nucleosomes and the interchromatin space to the partitioned NFRs. Adjacent NFRs form a platform favoring inter-NFR protein-mediated interactions. At a larger scale, multiple nanoscale domains coalesce forming higher-order structures that correspond to conventional TADs, potentially facilitated by cohesin-mediated loop extrusion.

Our novel insight is that this nuclear organisation can be generated purely through the biophysical properties of chromatin. We therefore predict that depletion of nucleosomes from a normally chromatinized region would be sufficient to drive it into the interchromatin space where it would become accessible to protein complexes including transcription factors (TFs) and the transcriptional machinery. Due to the difficulty of directly manipulating nucleosomes, we were only able to test this hypothesis *in silico*^14^. However, this idea is consistent with previous findings in yeast. For instance, nucleosome positioning—as determined by micrococcal nuclease sequencing—is sufficient to predict interaction domains in *S. cerevisiae*^18^. Moreover, yeast chromatin domains have been reconstituted *in vitro* from only DNA, histones, TFs and nucleosome remodelers^19^. Whether this principle extends to higher eukaryotes, where chromatin domains are approximately 100-fold larger, remains unclear^20^.

The histone chaperone Facilitates chromatin transcription (FACT) is a complex of two subunits: SSRP1 and SPT16. It was initially discovered as an accessory factor that allowed productive transcription of a chromatin template by the minimal transcriptional machinery *in vitro*^21^. Increasing evidence from *in vitro* and structural studies suggest that FACT engages with nucleosomes partially unraveled by the transcriptional machinery^22,23^, displaces histones by competitively occupying DNA-binding sites^24–26^ and transfers the H2A/B dimer upstream as RNAPII progresses through the gene, thereby simultaneously displacing histones and preventing their loss^27^.

The study of FACT *in vivo* has largely been limited to long-term perturbation experiments that have led to contradictory results regarding its role in transcription^28–37^ and nucleosome organisation^34–36,38–42^. Moreover, its role in 3D chromatin architecture has been minimally explored. Recent work has reported that acute degradation of FACT in human K562 cells leads to partial unwrapping of nucleosomes^43^ and minor changes in higher-order chromatin organisation, with TAD insulation subtly weakened^44^.

In this study we show that FACT plays a major role in both local and large-scale chromatin architecture using a rapid degradation system in mouse embryonic stem cells (mESCs), MCCu and a combination of genomic approaches. In the absence of FACT, nucleosomes around active promoters are depleted, leading to the loss of nanoscale domains and expansion of NFRs. The unprecedented resolution of MCCu reveals a rearrangement of chromatin interactions in the canonical promoter and newly accessible regions following FACT depletion. Surprisingly, and in contrast to previous studies, acute FACT degradation also leads to a dramatic increase in long-range promoter–promoter interactions that bypass TAD boundaries and can span up to tens of megabases. Moreover, we observe pervasive spreading of DNA binding proteins into the newly accessible DNA of active promoters. These results are consistent with a model in which nucleosome depletion is sufficient to drive genomic regions into the interchromatin space, where they are accessible to chromatin-binding proteins and can form long-range contacts with other promoters.

## RESULTS

### FACT loss leads to increased long-range promoter–promoter interactions that bypass TAD boundaries

To investigate the role of FACT in local chromatin structure, we generated a mESC cell line where the endogenous *Ssrp1* gene is tagged with an FKBP12^F36V^ degron domain^45^ (Figure 1A–B). Treatment with dTAG-13 resulted in decreased SSRP1 levels after 1h and virtually undetectable levels after 3h (as determined by immunoblotting and reference-normalised CUT&Tag) (Figure 1C–D, S1A). In addition, levels of SPT16—the other subunit of FACT—decreased despite not being tagged, which is consistent with previous work^36,43,46^(Figure 1E).

**Figure 1:**
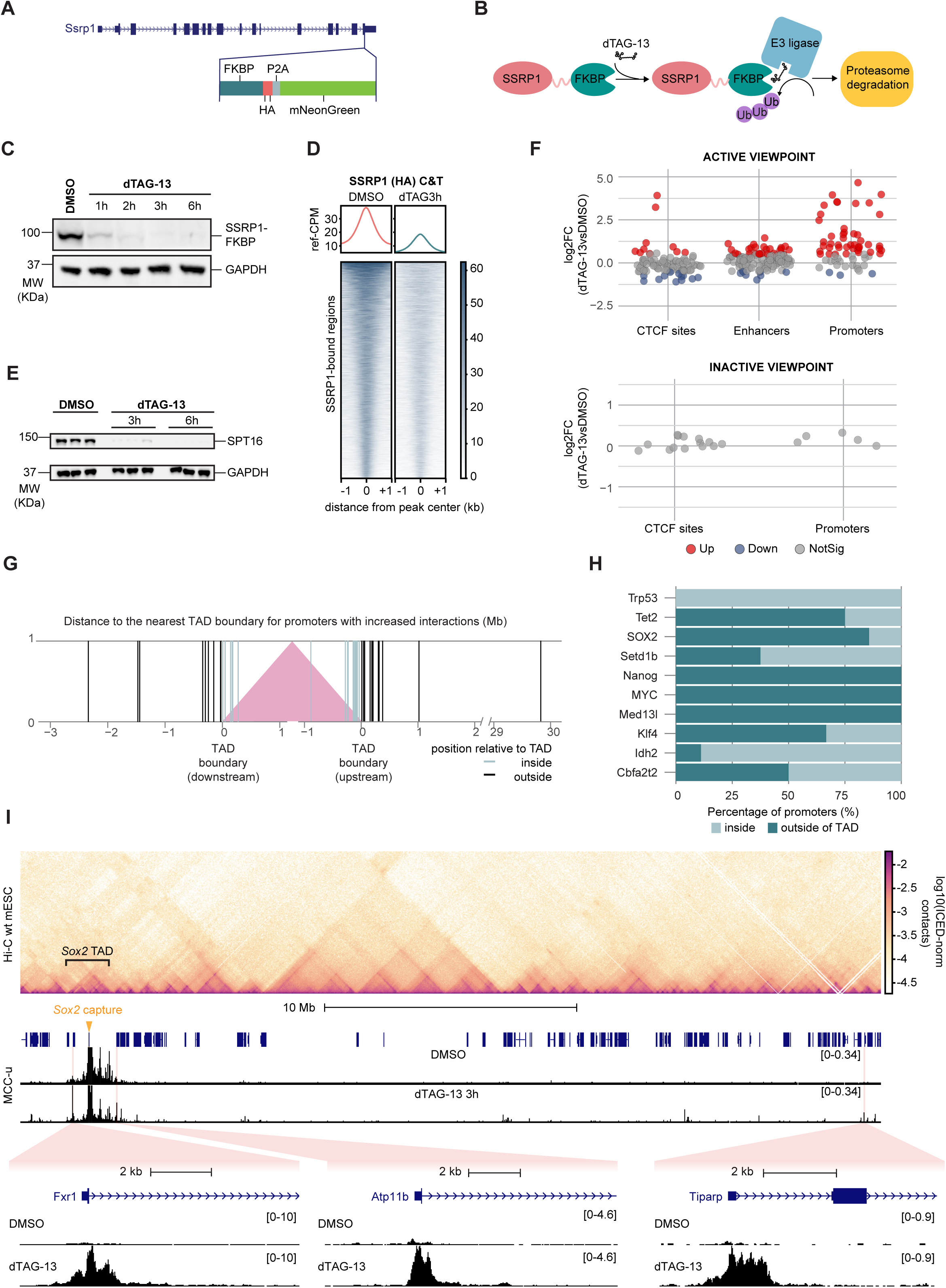
FACT depletion leads to increased interactions between promoters across topologically associating domains. (A) Schematic diagram of the addition of the FKBP degron domain to the endogenous *Ssrp1* locus. (B) Schematic representation of the degron system showing how SSRP1 is targeted for proteosomal degradation after dTAG-13 treatment. (C) Western blot analysis for SSRP1-FKBP (HA) in SSRP1-FKBP E14 cells treated with 1µM dTAG-13 for 1h, 2h, 3h, and 6h. Near-complete depletion is achieved after 3h. (D) Heatmap and mean distribution of SSRP1-FKBP (HA) CUT&Tag signal in cells treated with 1µM dTAG-13 for 3h showing a significant depletion in chromatin-bound SSRP1. (E) Western blot for SPT16 in SSRP1-FKBP E14 cells treated with 1µM dTAG-13 for 3h and 6h, showing that SPT16 is degraded despite not being tagged. (F) Statistical analysis of changes in MCCu interaction frequency after 3 h of dTAG-13 treatment between CTCF, enhancers, and promoters and transcriptionally active or inactive viewpoints. Active viewpoints include *Trp53, Sox2, Myc, Klf4, Tet2, Cbfa2t2, Med13l, Setd1b, Nanog,* and *Idh2*; inactive viewpoints include *Hba-a1* and *Hbb-b1*. Promoters were defined as ATAC-seq peaks within 1kb of a TSS (NCBI RefSeq), enhancers as ATAC-seq peaks overlapping H3K27ac peaks located >1kb from any TSS, and CTCF sites by CTCF ChIP-seq peaks. P values were calculated using the Wilcoxon rank-sum test and adjusted using the Benjamini–Hochberg method. (G) Distribution of distances between promoters with increased interactions upon FACT loss and the nearest TAD boundary of the capture viewpoint. Distances are shown separately for promoters whose closest boundary is the downstream or upstream boundary. TAD boundaries in mESCs were sourced from Bonev *et al.*^52^.(H) Proportion of promoters with increased interaction frequency within (light blue) and outside (teal) the corresponding TAD of each active viewpoint. Most capture viewpoints interact more frequently with promoters outside of their TAD. (I) Hi-C contact map for WT E14 cells (iterative correction and eigenvector decomposition [ICE] was used to normalise the matrices of read junction density) and MCCu signal in SSRP1-FKBP E14 cells around the *Sox2* locus (black tracks, CPT normalised to *cis*-interactions). The boundaries of the TAD comprising the *Sox2* locus have been indicated in the contact map. Examples of promoters with increased interactions with the *Sox2* promoter after 3h of treatment with 1µM dTAG-13 have been highlighted in pink and scaled to emphasize changes in interactions.

To test if the degron domain interfered with the function of FACT in gene expression we conducted RNA-seq. The results confirmed that WT and SSRP1-FKBP mESCs have a similar gene expression profile (Figure S1B–C). In line with previous work^36,47^, we also find that depletion of FACT leads to a dramatic decrease in cell number after 24h (Figure S1D). This shows that the complex is essential for mESC growth and highlights the importance of using a rapid degradation system to study it.

To investigate the effects of FACT loss on long-range chromatin interactions, we conducted MCCu for regions encompassing 10 promoters of actively transcribed genes and 2 promoters of inactive genes in SSPR1-FKBP mESCs. After 3h of dTAG-13 treatment, we detected significant changes in the frequency of *cis-*regulatory interactions with the active viewpoints, but not with the inactive ones (Figure 1F). This is consistent with FACT being recruited and functioning on actively transcribed genes^22,23^.

The most striking change following FACT depletion was that promoters gained significant numbers of additional long-range interactions with other promoters extending over distances up to 30 Mb, and both downstream and upstream of the corresponding gene (Figure S1E–F). Interestingly, these inter-promoter contacts crossed TAD boundaries, suggesting that FACT loss led to changes in higher-order chromatin organisation (Figure 1G–I, S1G). This is consistent with the observation by Mauksch *et al.* (2025)^44^ that TAD insulation is weakened by FACT loss in K562 cells.

### FACT depletion leads to the loss of nucleosomes and loss of nanoscale domains at promoters of active genes

To study the effects of FACT loss in local chromatin architecture, we generated multidimensional contact maps of promoters and proximal regions, which is possible since the tiling approach in MCCu captures contacts between all interacting fragments.

Since MCCu provides resolution at a subnucleosomal level, it reveals distinctive features undetectable with previous techniques. For instance, contact matrices display a striping pattern where the signal oscillates with a periodicity of 180-190bp. As previously shown, this pattern aligns with nucleosome positioning in a population of cells, where the position of each nucleosome may slightly vary but their spacing remains constant^14^ (Figure S2A).

Moreover, MCCu contact matrices at active promoters display interaction domains resembling TADs but 100- to 1000-fold smaller. These nanoscale domains correspond to nucleosome-containing regions where DNA linkers interact with each other and are partitioned by NFRs. In addition, NFRs interact with other NFRs (Figure S2B)^14^.

FACT depletion leads to the loss of the characteristic nucleosome-positioning pattern around promoters, as shown in of the *Sox2* and *Klf4* loci (Figure 2A–B, S2D). Notably, the extent of nucleosome perturbation decreases with increasing distance to the promoter (Figure S2D). To determine if nucleosome positioning changes upon FACT loss, we compared the distribution of distances between ligation junctions. Control samples have a multimodal distribution with a dominant mono-nucleosomal peak (∼190 bp). In contrast, FACT depletion leads to an increase in subnucleosomal ligation events (∼90 bp) and a reduction in mono- and di-nucleosomal ones (∼380 bp) (Figures 2C, S2E).

**Figure 2:**
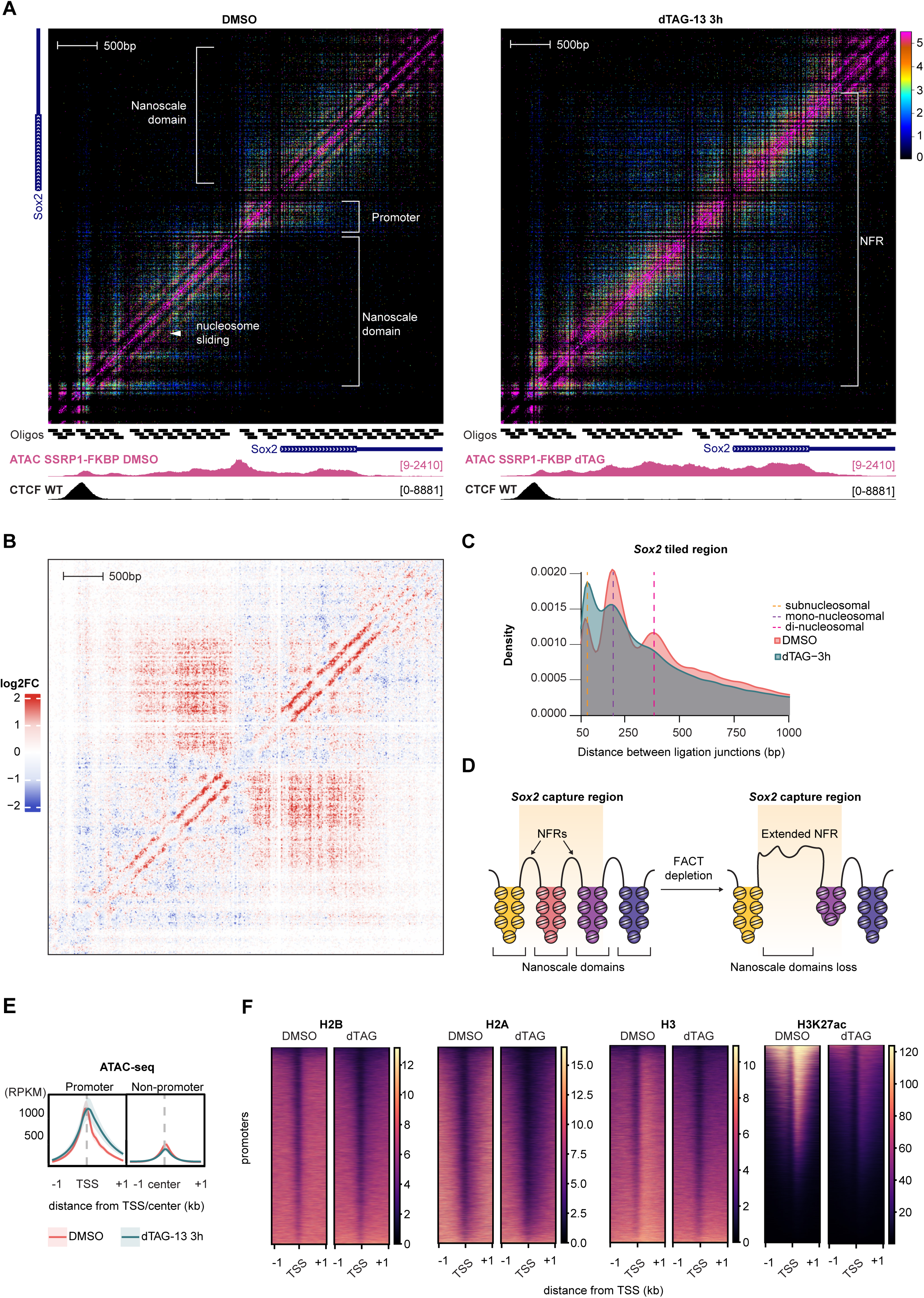
FACT depletion leads to nucleosome depletion and the loss of nanoscale domains at promoters of active genes. (A) Contact matrix (5bp resolution) at the *Sox2* promoter after treatment with DMSO, and dTAG-13 for 3h in SSRP1-FKBP E14 cells (ICE was used to normalise the matrices of read junction density; 6 replicates). Capture oligo positions, ATAC-seq (RPKM) for the DMSO control and dTAG-13 conditions are shown below the contact maps. CTCF ChIP-seq (CPM) from WT E14 is also shown. (B) Log2FC matrix upon FACT treatment shows the loss of nanoscale structures and the nucleosome stripping pattern. (C) Distribution of distances between ligation junctions in the tiled *Sox2* locus comparing DMSO (red) and 3h dTAG-13 (teal). (D) Model of nano-scale domains in the *Sox2* locus, showing nucleosome depletion and loss of nanoscale domains upon FACT loss. (E) Mean distribution of ATAC-seq signal (RPKM) at promoters and other accessible regions (non-promoters) after treatment with DMSO (red) and 1µM dTAG-13 for 3h (teal). Promoters were defined as region within 1kb a TSS (NCBI RefSeq) that overlap with ATAC-seq peak, and non-promoters as ATAC-seq peaks positioned >1kb from a TSS. Lines represent mean, shading represents ± SEM, *n* = 3 independent biological replicates (F) Heatmap representation of ChIP signal (ref-CPM) for H2B, H2A, H3 and H3K27ac at promoter regions after treatment with DMSO and 1µM dTAG-13 for 3h.

We further validated the nucleosome depletion upon FACT degradation with ATAC-seq and ChIP-seq for histone proteins. Consistently with MCCu results, FACT loss leads to an increase in chromatin accessibility around promoters, and a decrease in levels of H3, H2A, H2B, and H3K27ac (Figure 2E–F, S2C).

In the absence of FACT, we also observed the loss of nanoscale domains, whereby neighboring NFRs and affected nanoscale domains merge into a larger interaction domain. For instance, the two nanoscale domains flanking the *Sox2* promoter and the three nanoscale domains in the *Klf4* locus merge into a larger interaction domain upon FACT depletion (Figures 2A,B,D and S2D,F). By contrast, we do not detect any changes in the local chromatin environment of inactive genes upon FACT loss (Figure S2G–I), which is in line with FACT functioning on actively transcribed genes^48,49^.

Together, these findings suggest that FACT is required for maintaining intact nucleosomes and nano-scale domain architecture around the promoter region of actively transcribed genes.

### Increased promoter–promoter interactions are associated with spreading of chromatin–binding factors and increased chromatin accessibility

Next, we studied how chromatin-binding factors responded to the altered chromatin environment that followed FACT depletion. CUT&Tag profiling of transcription factors (TBP), co-activators (CBP, p300) and repressors (RING1B and KDM2B) showed that binding of all these factors increased and spread downstream of the promoter, whereas binding at other accessible regions remained largely unchanged (Figure 3A–B). As orthogonal validation, we carried out reference-normalised chromatin immunoprecipitation sequencing (ChIP-seq) for TBP, which also suggested increased protein binding at promoters and body of the gene (Figure S3A–B).

**Figure 3:**
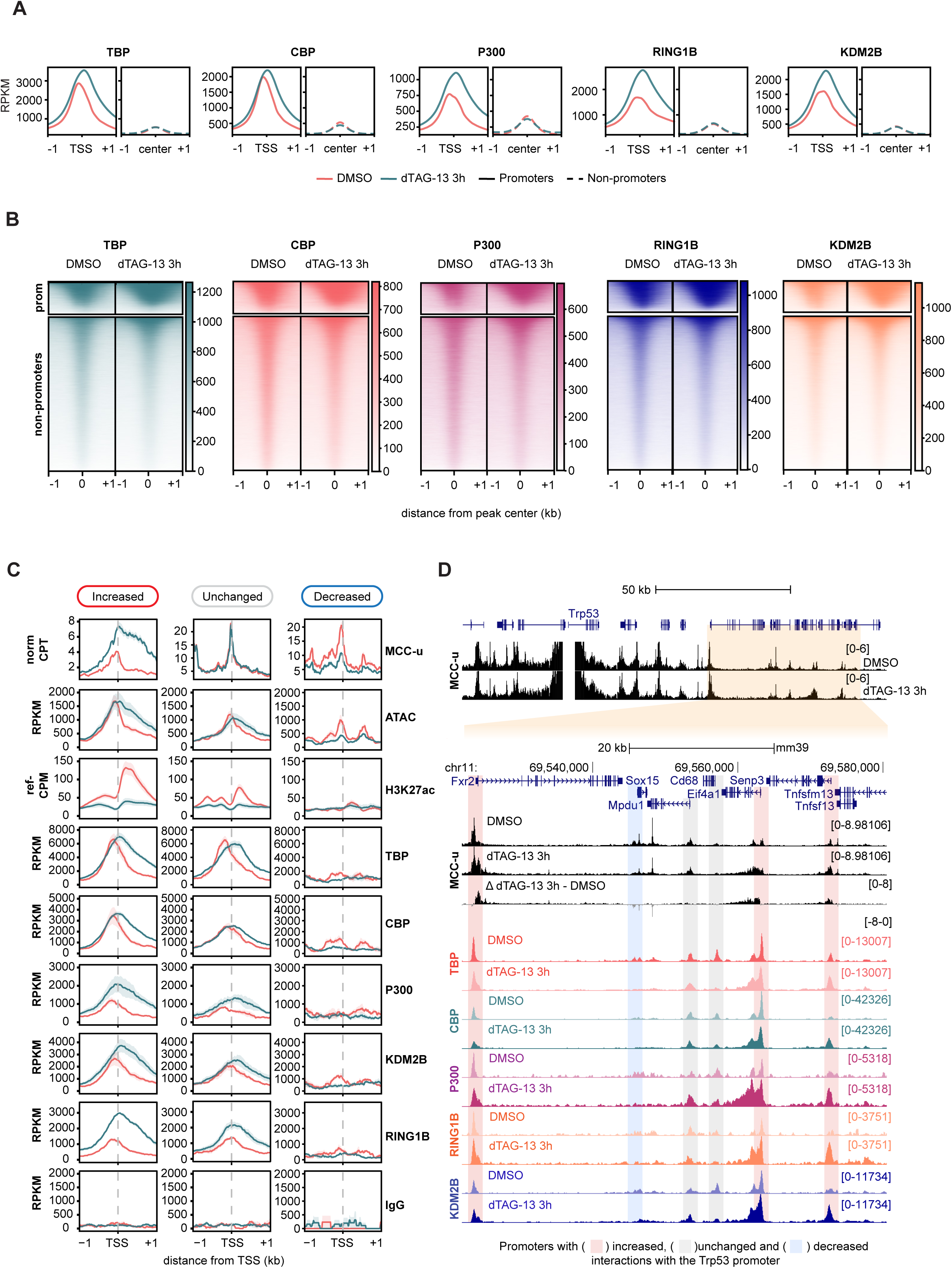
Spreading of protein complexes into the gene body upon FACT loss is associated with increased promoter–promoter interactions. (A) Mean distribution and (B) heatmap representation of CUT&Tag signal (RPKM) for TBP, CBP, P300, RING1B and KDM2B at promoters and other accessible regions (non-promoters) after treatment with DMSO (red) and 1µM dTAG-13 for 3h (teal). Promoters were defined as region within 1kb a TSS (NCBI RefSeq) that overlap with ATAC-seq peak, and non-promoters as ATAC-seq peaks positioned >1kb from a TSS. (C) Mean distribution of signal for MCCu (*cis*-norm-CPT), ATAC (RPKM), H3K27ac (ref-CPM), TBP, CBP, p300, KDM2B, RING1B, and IgG CUT&Tag (RPKM) at promoters with increased, unchanged and decreased interactions. Lines represent mean, shading represents ± SEM (*n* = 3 independent replicates for all assays and *n* = 6 for MCCu) (G) MCCu and CUT&Tag signal (RPKM) for TBP, CBP, p300, KDM2B and RING1B at genes that interact with the *Trp53* promoter.

Next, we studied how the changes in protein binding, chromatin accessibility, and H3K27ac levels associated with changes in promoter–promoter interactions after FACT depletion (Figure 3C–D). The results show a strong association between protein spreading, increased chromatin accessibility and increased promoter–promoter interactions. After FACT loss, protein binding and ATAC signal increased dramatically at promoters with increased interactions, whereas the changes were more subtle at other promoters. However, H3K27ac levels dropped across all promoters regardless of changes in contact frequency.

Overall, the results suggest that the decrease in H3K27ac does not contribute to the changes in promoter–promoter interactions upon FACT loss—although this cannot be extended to other acetylation marks we have not tested. Instead, the findings point to nucleosome loss and increased spreading of protein complexes as potential drivers of increased promoter interactions.

### Subnucleosomal chromatin interactions at canonical promoters are rewired upon FACT loss

To visualize contacts between DNA-bound factors within promoters, we performed contact sequence reconstruction analysis with the MCCu data, as previously described^14^. In short, the principle of this approach is that DNA binding proteins protect the sequence from MNase digestion. Thus, we can determine whether a chromatin-bound protein is upstream or downstream of the ligation junction. The directionality of the reads can be used to pinpoint exact binding positions and interactions between DNA-associated proteins.

Our analysis shows that there is a rearrangement in chromatin interactions at canonical promoters following FACT loss. As shown in the *Setd1b* and *Klf4* the promoters (Figure 4A–B), the punctate interactions between factors that are detected at canonical promoter regions decrease upon FACT depletion and contacts are gained at the newly nucleosome-depleted regions. Interestingly, we observe spreading of proteins (e.g. TBP) into the regions that gain interactions upon FACT loss. This suggests that aberrant binding of protein complexes to the newly accessible regions leads to the rearrangement of local chromatin interactions in the promoter.

**Figure 4:**
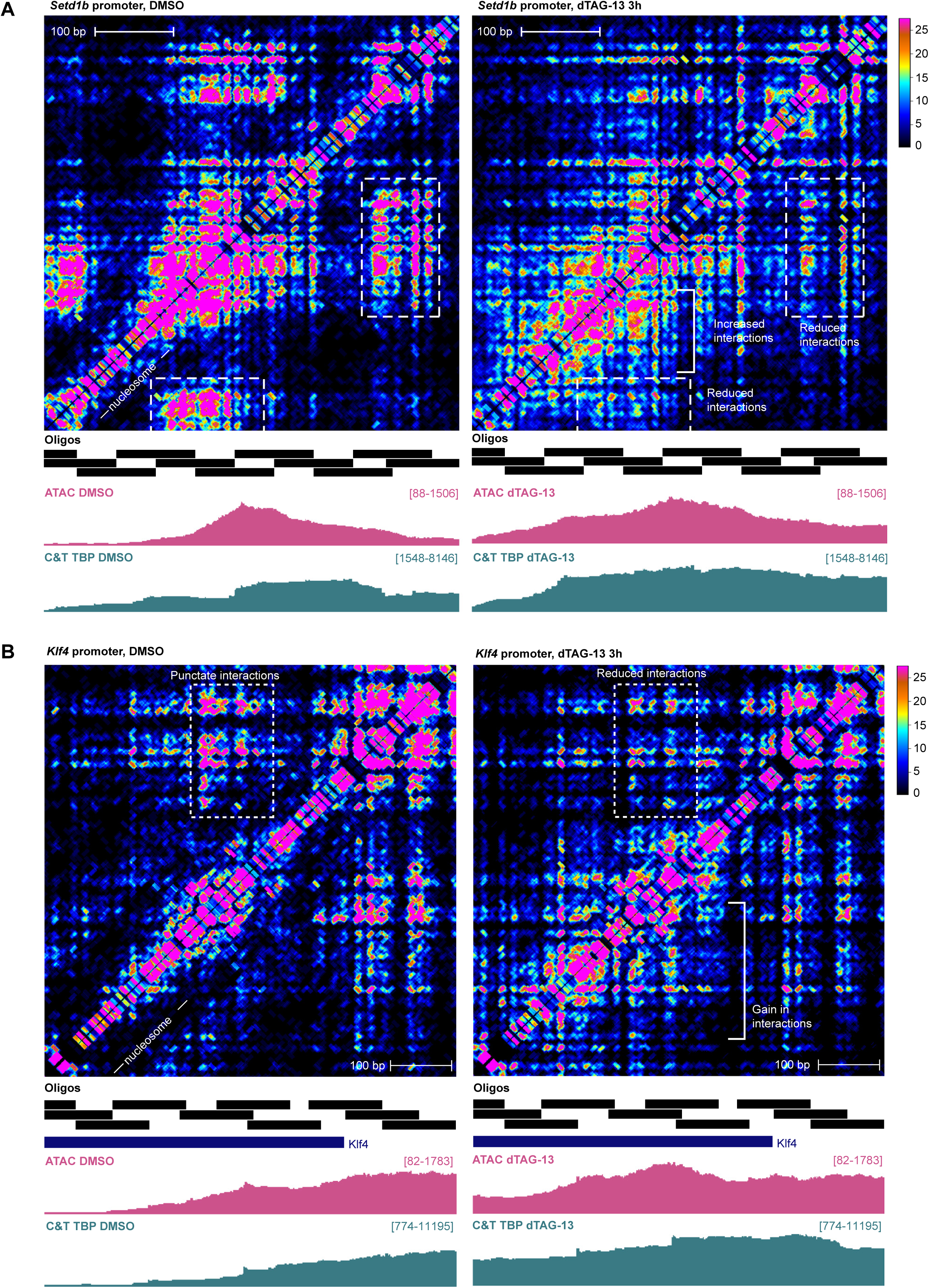
FACT loss leads to the rearrangement of subnucleosomal chromatin interactions at canonical promoters. (A) Contact sequence reconstruction map (1bp resolution) at the *Setd1b* promoter (chr5:123278895–123279525) and (B) the *Klf4* promoter (chr4:55531978–55532778) Contact matrices were normalised to the total number of interactions (n=6 replicates). Capture probe positions, ATAC-seq (RPKM) and, CUT&Tag for TBP (RPKM) for the control and treatment conditions are shown below the contact maps. Changes in interactions have been indicated.

### Disruption in chromatin organisation is more widespread than transcriptional changes upon FACT loss

We investigated the effects that FACT loss had on transcription by carrying out reference-normalised transient transcriptome sequencing (TT-seq). FACT degradation primarily led to the downregulation of very highly expressed genes (Figure 5A–C), supporting the idea that the transcriptional machinery perturbs chromatin structure and that FACT is key for its maintenance^22,23^.

**Figure 5:**
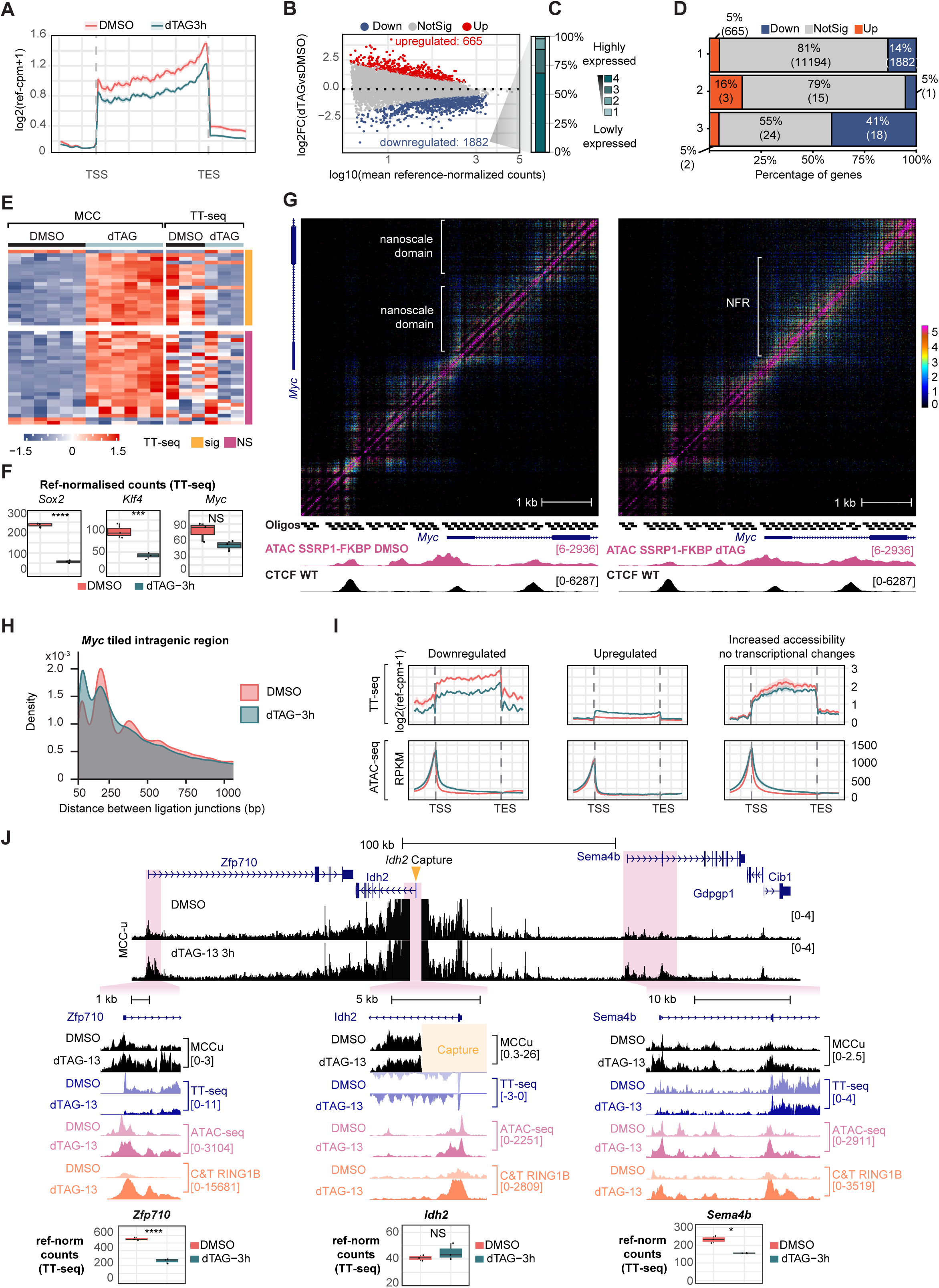
Disruption in chromatin organisation is more widespread than transcriptional changes upon FACT loss. (A) Mean distribution of TT-seq signal (ref-CPM) at protein-coding genes in SSRP1-FKBP E14 cells treated with DMSO (red) and dTAG-3h for 3h (teal). Lines represent mean, shading represents ± SEM, *n* = 3 independent biological replicates. (B) MA plot of TT-seq data showing the effect of 3h of dTAG-13 treatment in SSRP1-FKBP E14 cells in nascent transcription (C) Distribution of downregulated genes upon dTAG-13 treatment based on expression levels. (D) Proportion of differentially expressed genes following FACT degradation across (1) the genome, (2) genes with unchanged or decreased promoter–promoter interactions, and (3) genes with increased promoter–promoter interactions. (E) Heatmap comparing promoter–promoter interactions (MCCu) and nascent transcription (TT-seq) (z-score). Rows have been clustered based on statistical significance of transcriptional changes (TT-seq). (F) Reference-normalised TT-seq counts corresponding to the *Sox2*, *Klf4* and *Myc* in the DMSO control (red) and dTAG-13 3h treatment samples (teal). P-values were determined with differential gene expression analysis with DESeq2 (**** <1e−4, *** <1e−3, ** <1e−2, * <5e−2, NS >5e−2) (G) Contact matrix (5bp resolution) at the *Myc* promoter after DMSO and 3h of dTAG-13 treatment in SSRP1-FKBP E14s. (ICE was used to normalise the matrices of read junction density; 6 replicates). Capture oligo positions, ATAC-seq (RPKM) for the DMSO control and dTAG-13 conditions are shown below the contact maps. CTCF ChIP-seq (CPM) from WT E14 is also shown. (H) Distribution of distances between ligation junctions in the tiled intragenic region of *Myc* comparing DMSO (red) and 3h dTAG-13 (teal). (I) Mean distribution of TT-seq (ref-CPM) and ATAC-seq signal (RPKM) across gene bodies for downregulated genes, upregulated genes and genes with increased accessibility but no significant transcriptional changes following FACT depletion. Lines represent mean, shading represents ± SEM, *n* = 3 independent biological replicates. (J) MCCu (black, *cis*-norm CPT), TT-seq (blue, ref-CPM), ATAC-seq (pink, RPKM) and CUT&Tag signal for RING1B (orange, RPKM) at the *Idh2* locus and surrounding genes in the DMSO control and 3h dTAG-13 treatment sample. Magnified views of the highlighted regions are provided to emphasize the changes. Reference-normalised TT-seq counts corresponding to the highlighted genes are indicated below.

We observe a significant association between increased promoter–promoter interactions and transcriptional downregulation (Fisher’s exact test*, p-value*=0.005) (Figure 5D). However, increased promoter–promoter interactions are not always associated with a decrease in transcription (Figure 5E).

Changes in local chromatin structure are also not consistently associated with a transcriptional output. In some cases, the loss of nanoscale domains coincided with transcriptional downregulation (e.g. *Sox2* and *Klf4*, Figure 2A–D, S2D–F, 5F). In contrast, other loci showed no significant changes in transcription, despite alterations in overall domain structure (e.g. *Myc* and *Idh2*, Figure 5F–H, S4A–B).

Overall, alterations in chromatin structure were more widespread than transcriptional changes upon FACT degradation—a subset of genes displays increased chromatin accessibility yet without significant transcriptional changes (Figure 5I–J). These results suggest that FACT primarily has a role in maintaining chromatin architecture, and that breakdown of higher-order chromatin organisation does not necessarily result in transcriptional changes. However, if there is a transcriptional change, the breakdown in chromatin organisation generally results in decreased transcription.

## DISCUSSION

In this study we show that FACT depletion leads to dramatic changes in the structure of promoters through loss of local nucleosomes, which results in large-scale and local disruption of chromatin architecture. Moreover, by integrating different genomic approaches, we have provided a detailed insight into the dysregulation of chromatin factor binding and transcription that follows FACT depletion.

Our results redefine the role of FACT as a major regulator of chromatin structure. We show that FACT is key for nucleosome maintenance *in vivo* since its depletion leads to nucleosome loss around the promoters of actively transcribed genes. This differs from the results recently reported by Žumer *et al* (2024), where they only detect an increase in partially unwrapped nucleosome intermediates (potentially hexasomes) after rapid FACT degradation in K562 cells. However, it has been shown that multiple rounds of transcription of a chromatinised template in the absence of histone chaperones leads to the formation of hexasomes and, subsequently, nucleosome loss^50^. Thus, this disagreement could potentially be explained by differences in transcriptional activity and degradation dynamics between the cell lines^43^—it has been suggested that embryonic stem cells are particularly transcriptionally active^51^.

Moreover, we demonstrate that FACT is important for orchestrating chromatin architecture at promoters. After FACT degradation, nucleosome depletion around promoters leads to the loss of nanoscale domains and the emergence of newly accessible regions that coalesce with neighbouring NFRs. Thus, FACT loss leads to the formation of larger NFRs that favour binding of protein complexes and have increased long-range interactions with other promoters. The observation that these new 3C contacts can bypass TAD boundaries is consistent with previous findings showing that TAD insulation is weakened in the absence of FACT^44^. In addition, the stronger changes in chromatin structure we observe relative to previous work^43,44^ may result from more rapid and extensive chromatin disruption in mESCs and from the inability of previous 3C techniques to resolve nanoscale structures within genic regions, where FACT primarily functions.

From a broader perspective, this degradation system provides *in vivo* evidence for a model of chromatin organisation we recently proposed. In this model, chromatin folds into nanoscale domains that are partitioned by NFRs, which interact with adjacent NFRs in the interchromatin compartment—the space in the nucleus between chromatin domains that is enriched in proteins and RNAs as shown by super-resolution imaging. We propose that chromatin folding is largely driven by the intrinsic properties of nucleosomes; however, this hypothesis has only been tested *in silico* due to the technical difficulties associated to directly manipulating nucleosomes^14^.

Acute degradation of FACT indirectly shows that localised nucleosome loss leads to the rearrangement of nanoscale domains, supporting a central role for nucleosomes in chromatin architecture. Moreover, these newly accessible regions exhibit an increase in protein binding, which aligns with their prediction that NFRs coalesce above aggregated nucleosomes into the interchromatin compartment (Figure 6A–B). Overall, although direct manipulation of nucleosomes would be necessary for definitive proof, such approaches remain technically challenging, making our system a close experimental approximation of this ideal.

**Figure 6:**
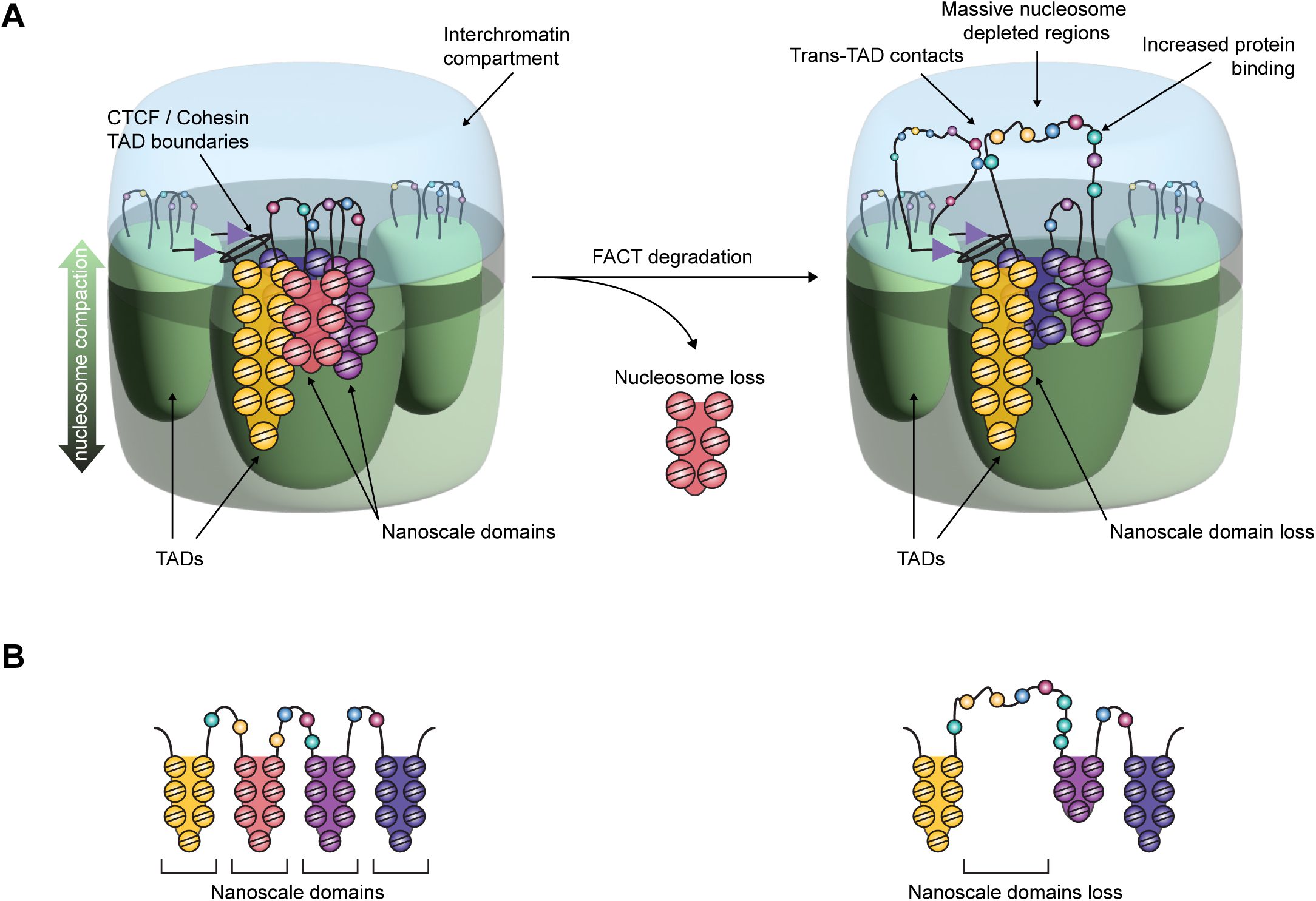
Model summary of the effect of FACT depletion on chromatin structure. (A) Three-dimensional architectural model of nanoscale domains within TADs following FACT depletion. In the absence of FACT, nucleosomes are depleted at promoters leading to the expansion of nucleosome-free regions. These enlarged NFRs coalesce above aggregated nucleosomes into the interchromatin compartment, where they are accessible for binding of protein complexes. These extended NFRs can bypass TAD boundaries to interact with other promoters. (B) Schematic representation of the linear genome folding corresponding to the three-dimensional model in (A).

In summary, by combining a high-resolution tiled 3C technique with genomic approaches we have provided detail insights into the effects of FACT in chromatin structure and function and, thus, demonstrated that it plays a major role in genome architecture *in vivo*. Beyond advancing our understanding of FACT, this degradation system has served as an indirect approach to deplete nucleosomes *in vivo* and supports an emerging model that positions nucleosomes and their intrinsic biophysical properties as central drivers of chromatin structure.

## LIMITATIONS OF THE STUDY

We generated a degron system to rapidly degrade SSRP1 in mESCs and minimise compensatory mechanisms; however, we cannot completely rule out secondary effects on chromatin structure and transcription. Furthermore, the conclusions on the role of FACT in 3D chromatin architecture are based on the data captured for 12 gene promoters, which only represent a fraction of the genome and might not be fully generalisable across all genes. We conclude that FACT opposes to protein spreading based on CUT&Tag and ChIP-seq for 5 chromatin-binding factors. Although these experiments provide us with a valuable mechanistic insight, we cannot confirm that all chromatin-binding factors respond the same way. Finally, in most assays we used ectopic spike-in for reference normalisation, but we cannot exclude minor normalisation biases due to cell counting or pipetting errors.

## RESOURCE AVAILABILITY

Requests for further information and resources should be directed to and will be fulfilled by the lead contact Thomas Milne

Weatherall Institute of Molecular Medicine, University of Oxford, Oxford OX3 9DS, UK

### Materials availability

All unique/stable reagents generated in this study are available from the lead contact with a completed materials transfer agreement.

### Data and code availability

The code for analysis of MCCu data is available for academic use through the Oxford University Innovation software store (https://process.innovation.ox.ac.uk/software/p/16529a/micro-capture-c-academic/1). Other code is available through GitHub (https://github.com/Davies-Genomics-and-Genome-Editing-Lab/MCCu). The SeqNado pipeline used for the analysis of polyA RNA-seq, ATAC-seq, ChIP-seq, CUT&Tag and TT-seq is available in https://pypi.org/project/seqnado.

Raw high-throughput sequencing data and processed files generated in this study have been deposited in the NCBI Gene Expression Omnibus (GEO) under the accession number GSE314917 (polyA RNA-seq, MCCu, ATAC-seq, ChIP-seq, CUT&Tag, TT-seq). Previously published data reused in this study can are available under the following GEO accession numbers: GSE96107 (Hi-C data for wildtype E14s) and GSE123670 (ChIP-seq data for CTCF in wildtype E14s).

## ACKNOWLEDGMENTS

The authors acknowledge the flow cytometry facility at the Medical Research Council (MRC) WIMM for providing assistance with cell sorting and the Genome Engineering & Transgenics Facility Team for sharing their E14 Tg2a IV cells with us. Thanks to R.J. Klose, E. Dimitrova and H.Y.A. Au for sharing the anti-KDM2B antibody and *Drosophila melanogaster* S2 cells with us.

T.A.M., and A.M.D.-F. were funded by Medical Research Council (MRC, UK) Molecular Haematology Unit grant MC_UU_00029/6. R. A. B. was supported by Wellcome (224135/Z/21/Z). C.C. was supported by a Wellcome Trust Genome Medicine and Statistics studentship. J.L.T.D. was supported by the Keble College Sloane Robinson Scholarship, the Clarendon Fund, and a Radcliffe Department of Medicine Studentship. J.O.J.D., J.L.T.D., and H.L. were supported by the Lister Institute, the MRC Molecular Haematology Unit (MC_UU_00029/04), and Wellcome (225220/Z/22/Z).

## AUTHOR CONTRIBUTIONS

T.A.M., R.A.B., and A.M.D.-F. conceived the project and designed experiments. A.M.D.-F. generated the degron cell line and carried out experiments. A.M.D.-F., C.C., H.L. and J.L.T.D. analysed and curated the data. T.A.M., R.A.B., J.O.J.D., and A.M.D.-F. interpreted the data and wrote the manuscript. T.A.M., R.A.B., and J.O.J.D. provided supervision. T.A.M. provided funding. All authors reviewed the manuscript.

## STAR★METHODS

### KEY RESOURCES TABLE

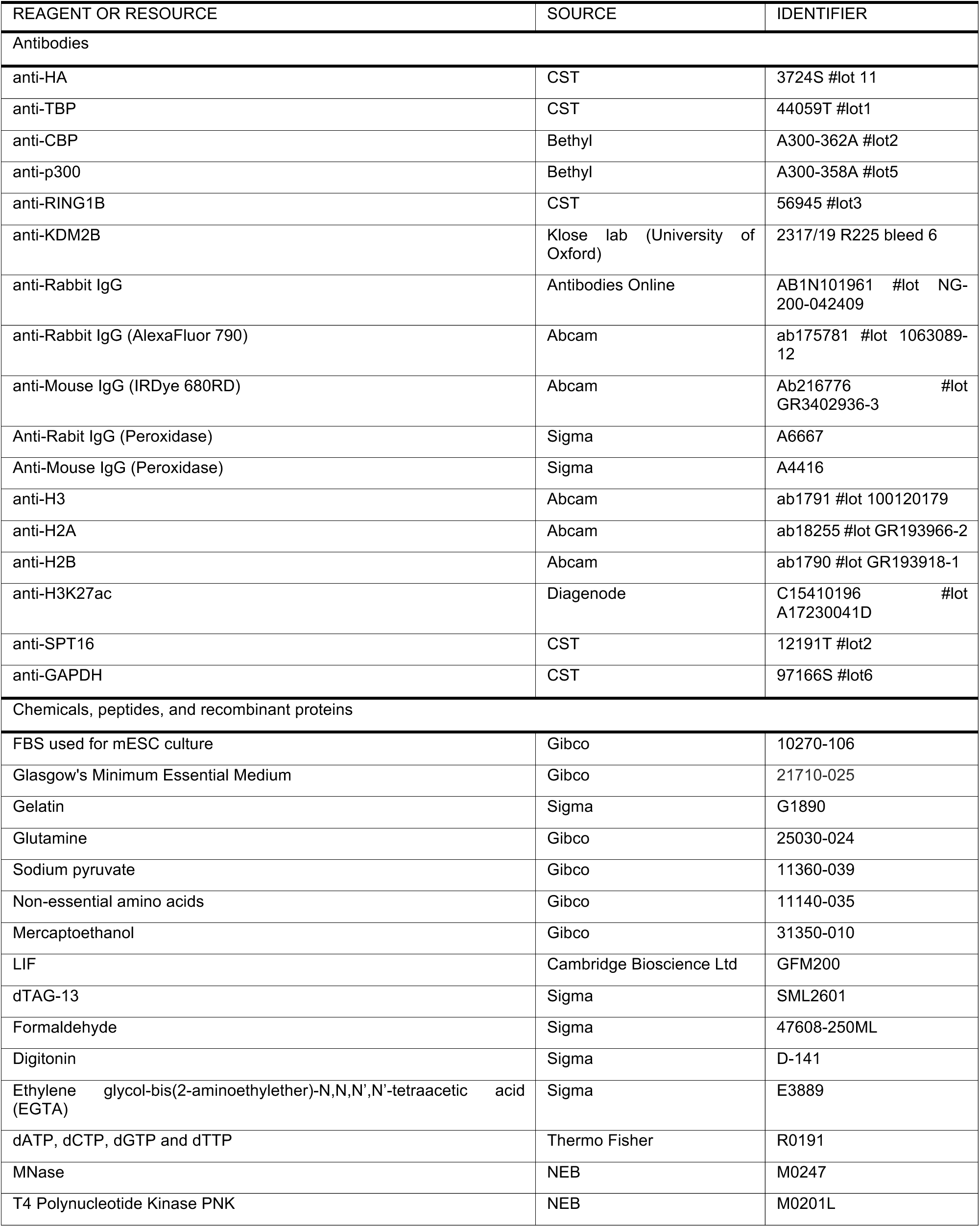
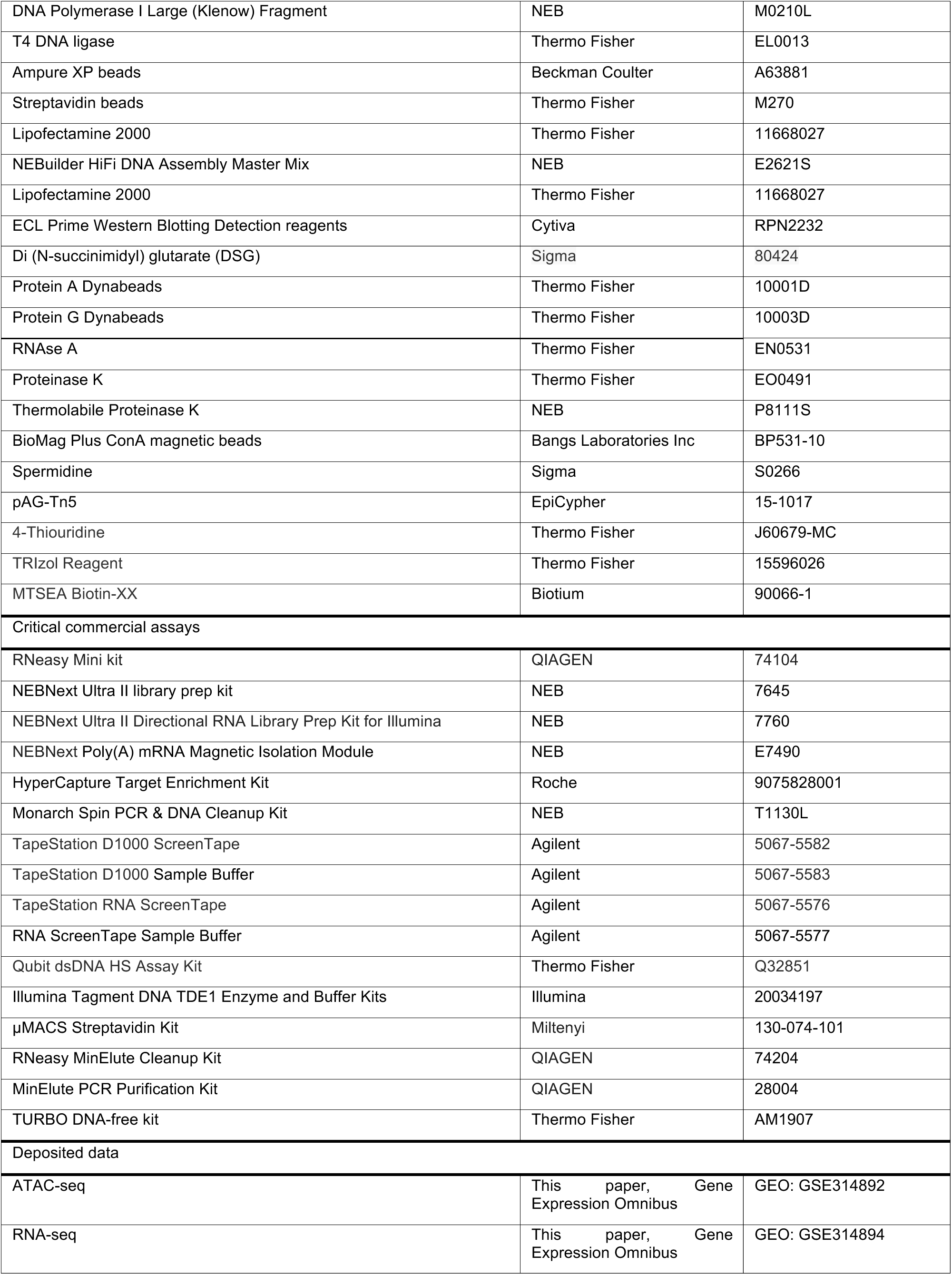
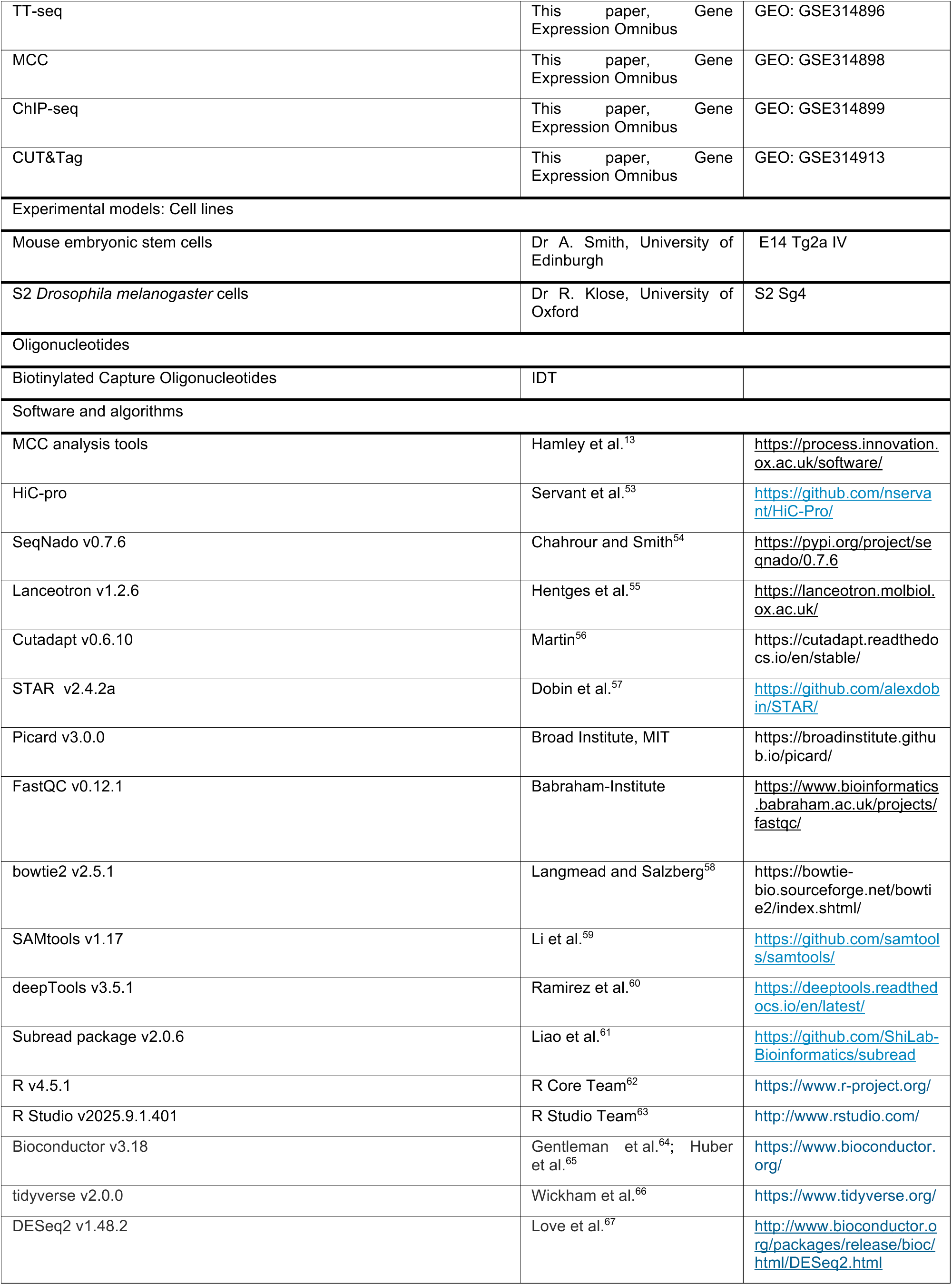
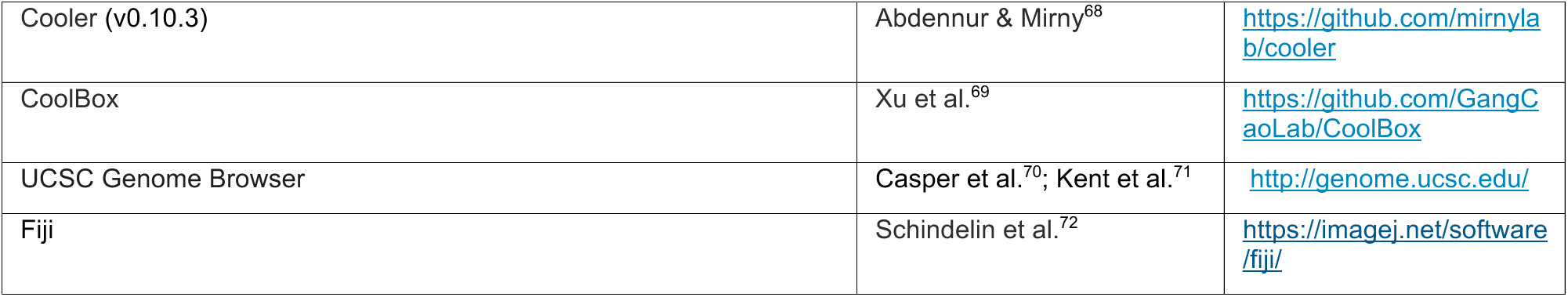

### CELL LINES

The murine embryonic stem cell line (E14 Tg2a IV, male biological origin) was grown in plates coated with 0.1% gelatin (Sigma, G1890) with Glasgow MEM x1 liquid (Gibco 21710-025) complemented with 10% FCS (Gibco 10270-106), 1x Glutamine (Gibco 25030-024), 1 mM sodium pyruvate (Gibco 11360-039), 1x non-essential amino acids (Gibco 11140-035), 100 uM mercaptoethanol (Gibco 31350-010) and 10 ng/mL LIF(Cambridge Bioscience Ltd GFM200). Cells were grown in a humidified incubator at 37*°*C with 5% CO_2_.

## METHOD DETAILS

### GENERATION OF DEGRON CELL LINES

By means of CRISPR/Cas9-mediated homology-directed repair (as previously described^37^), FKBP12^F36V^-HA-HA-P2A-mNeonGreen was added before the stop codon in both *Ssrp1* alleles of E14 cells. Using Lipofectamine 2000, cells were transfected with pX458 (encoding Cas9, sgRNA targeting the C-terminus of *Ssrp1* and the fluorophore mRuby) and an HDR plasmid (containing the FKBP12^F36V^-HA-HA-P2A-mNeonGreen sequenced flanked by 500bp homology arms to target the end of the *Ssrp1* gene. After 24h, mRuby-positive cells were isolated and grown for 2 weeks. Then, mNeonGreen positive-cells isolated and grown as colonies to select individual clones. Screening for homozygous clones with the insertion was carried out by western blotting.

### GROWTH ASSAY

SSRP1-FKBP E14 cells were treated with DMSO and dTAG-13 as described above for 3h, 6h, 24h and 48h. Cells attached to the plates were washed with PBS and fixed and permeabilised with methanol:acetone (1:1). Trypan blue diluted 2x in PBS was used to stain the plates. Residual dye was removed by carrying out with three PBS washes.

### WESTERN BLOTTING

Proteins were extracted from 1×10^6^ cells using a high-salt buffer (20 mM Tris-HCl pH 8.0, 300 mM KCl, 5 mM EDTA, 20% glycerol, 0.5% IGEPAL CA-630, protease inhibitor cocktail). Proteins were denatured at 95°C, separated using NuPAGE Bis-Tris 4–12% gels and transferred to polyvinylidene fluoride membranes. Antibodies used are listed in the key resources table.

#### ChIP-seq

Chromatin immunoprecipitation was carried out as previously described^73^. In short, 5×10^6^ cells were single-fixed (1% formaldehyde for 10min) for histones and histones modifications. For transcription/chromatin factors, cells were double-fixed (2mM disuccinimidyl glutarate for 30 min, then 1% formaldehyde for 30 min) and lysed in 120uL SDS lysis buffer (10 mM Tris-HCl pH 8.0, 1 mM EDTA, 1% SDS). The lysate was then syringe-passaged through a 27-gauge needle to enhancer the lysis. To generate fragments between 200-300bp, chromatin was sonicated with a Covaris ME220. Soluble chromatin was isolated by maximum speed centrifugation, diluted 10x (0.01% SDS, 1.1% Triton-X, 1.2mM EDTA, 16.7mM Tris-HCl, 167mM NaCl) and chromatin from *Drosophila* S2 cells was spiked in a ratio of 1:4 (S2:E14). To reduce non-specific binding, chromatin was incubated with 5uL of A- and G-coupled dynabeads (Thermo Fisher 10001D, 10003D) for 15min at 4°C with rotation. Then, 20% of the sample was taken as input and the rest was incubated overnight with 2uL of the corresponding antibody (see key resources table). To isolate chromatin-antibody complexes, 15uL of Protein A- and G-coupled dynabeads were used, followed by 3 washes with RIPA buffer (50 mM HEPES-KOH pH 7.6, 500 mM LiCl, 1 mM EDTA, 1% NP40 and 0.7% sodium deoxycholate) and once with 50mM NaCl Tris-EDTA buffer. Samples were eluted from beads with SDS lysis buffer. Then, they were treated with RNase A, Proteinase K and decrosslinked by incubation at 65°C overnight. DNA was purified with the Monarch Spin PCR & DNA Cleanup Kit (NEB T1130L) and sequencing libraries were generated with the NEBNext Ultra II library preparation kit (NEB 7645). Libraries were sequenced by 150 bp paired-end sequencing with a NovaSeq X Plus (Illumina).

#### ATAC-seq

ATAC-seq was carried out as previously described^74^. Fifty thousand cells were harvested, washed with cold PBS and nuclei were extracted by incubation with 50uL cold lysis buffer for 3 minutes on ice (10mM Tris-HCl, 10mM NaCl, 3mM MgCl2, 1% NP-40, 1% Tween-20 and 0.5% Digitonin). Then, 1mL of wash buffer (10mM Tris-HCl, 10mM NaCl, 3mM MgCl2, 1% Tween-20) was added to the lysate. Nuclei were pelleted and resuspended in a transposase reaction mix (Illumina 20034197) at 37°C for 30 minutes. Next, DNA was purified with the MinElute column kit (Qiagen 28004). DNA was then amplified for 10 PCR cycles and purified again with the MinElute kit. Samples were sequenced by 150 bp paired-end sequencing with a NovaSeq X Plus (Illumina).

#### CUT&Tag

CUT&Tag was carried out as previously described^75^. Briefly, nuclei from 25,000 cells were extracted and permeabilised by incubating the cells with lysis buffer (20mM HEPES-KOH, 10mM KCl, 0.5mM spermidine, 1% (v/v) Triton X, 20% (v/v) glycerol, 1:200 proteinase inhibitor cocktail) at 4°C for 10 minutes. Nuclei were lightly fixed with 0.1% formaldehyde for 2 minutes. To prevent overfixation, the reaction was quenched with 75mM Glycine. Nuclei were pelleted and resuspended in wash buffer (20mM HEPES-KOH, 150mM NaCl, 0.5mM spermidine). Then, 5uL of activated BioMag Plus ConA magnetic beads (20mM HEPES-KOH, 10mM KCl, 1mM CaCl_2_, 1mM MnCl_2_) were added and incubated for 5 minutes at RT to allow binding of nuclei to beads. The wash buffer was replaced with 25uL antibody buffer (0.1% BSA, 2mM EDTA, 20mM HEPES-KOH, 150mM NaCl, 0.5mM spermidine) and the nuclei were incubated with 0.5uL of the primary antibody for 1h at RT (see key resources table). Then, nuclei bound to the magnetic beads were resuspended in 25uL of fresh antibody buffer with secondary antibody (1:100) and incubated at RT for 30 minutes. After the antibody incubation, beads were washed with 150uL of wash buffer. The beads were resuspended in 25 uL wash buffer with 1:20 pAG-Tn5 (EpiCypher 15-1017) and incubated for 1h at RT to allow pAG-Tn5 binding. Samples were washed twice with a high-salt wash buffer (300 mM NaCl, 20mM HEPES-KOH, 0.5mM spermidine) to reduce non-specific binding. The transposition reaction was carried out at 37°C for 1h in tagmentation buffer (10mM MgCl_2_, 300 mM NaCl, 20mM HEPES-KOH, 0.5mM spermidine). Next, the samples were washed with TAPS wash buffer (10mM TAPS, 200uM EDTA). For particle release, the samples were resuspended in 5uL of SDS buffer with thermolabile Proteinase K (NEB P8111S) (1% SDS, 10mM TAPS, 10% (v/v) ProtK) and incubated at 37°C for 1h and 58°C for 1h. 15uL of Triton neutralization solution was added to the samples. DNA was amplified for 14 cycles with NEXTera indices and NEBNext Ultra II Q5 Master mix. DNA was purification was carried out with AMPure XP beads (Beckman Coulter A63881). Samples were sequenced by 150 bp paired-end sequencing with a NovaSeq X Plus (Illumina).

#### TT-seq

TT-seq was conducted as previously described. In short, 3×10^7^ E14 cells were treated with 500uM 4sU (Thermo Fisher J60679-MC) for 5 minutes before RNA was isolated by Trizol extraction. For reference normalisation, S2 *Drosophila* cells were also treated with 500uM 4-thiouridine, lysed in Trizol and added to the samples in a 1:2 ratio (S2:E14). To remove residual DNA, samples were treated with the TURBO DNA-free kit and RNA quality was assessed with TapeStation. Total RNA was fragmented by sonication (Covaris ME220), 4sU labelled RNA was biotinylated with (Biotium 90066-1) and pulled down with the µMACS Streptavidin Kit (Miltenyi 130-074-101). Isolated RNA was purified with the RNeasy MinElute Cleanup Kit (Qiagen 74204). Sequencing libraries were generated with NEBNext Ultra II Directional RNA Library Preparation Kit for Illumina (NEB 7760) and sequenced by 150 bp paired-end sequencing using a NovaSeq X Plus (Illumina).

#### PolyA RNA-seq

One million E14 cells were collected and washed with PBS. For reference normalisation purposes, *Drosophila* S2 cells were added in a ratio of 1:2 (S2:E14). Total RNA was extracted with the RNeasy Mini kit (Qiagen 74104) and RNA quality was assessed with TapeStation. PolyA mRNA was isolated with the NEBNext Poly(A) mRNA magnetic isolation module (NEB E7490) as indicated by the manufacturer. Then, sequencing libraries were generated with the NEBNext Ultra II Directional RNA Library Prep Kit for Illumina (NEB 7760). Samples were sequenced by 150 bp paired-end sequencing with a NovaSeq X Plus (Illumina).

### MCCu LIBRARY GENERATION AND CAPTURE

MCCu was conducted as previously described^13,14^. In short, 10^7^ cells were fixed with 2% formaldehyde (Sigma, 47608-250ML) for 10 minutes at room temperature (RT). The reaction was quenched with 130mM glycine. The cells were pelleted, washed with PBS and permeabilize with 0.005% digitonin for 15min. The cells were resuspended in reduced-calcium MNase buffer (10 mM Tris-HCl pH 7.5, 1 mM CaCl_2_) and split into three aliquots. Each aliquot was treated with a titration of different micrococcal nuclease (NEB) concentrations (8-10 Kunitz U) for 1h at 37°C and the reaction was stopped by adding ethylene glycol-bis(2-aminoethylether)-N,N,N′,N′-tetraacetic acid (EGTA) at 5mM. After a PBS wash, the cells were resuspended in DNA ligase buffer (Thermo Fisher EL0013) and incubated with 200u/mL of T4 Polynucleotide Kinase PNK (NEB M0201L), 100U/mL of DNA Polymerase I Large (Klenow) Fragment (NEB M0210L) and 300 U/mL of T4 DNA ligase (Thermo Scientific EL0013) for 2h at 37°C and 8h at 20°C. Chromatin crosslinking was reversed by incubation at 65°C with Proteinase K and DNA was purified with phenol-chloroform extraction. Digestion and ligation efficiencies were assessed by TapeStation (Agilent D1000). The two best ligated samples for each biological replicate were selected for sonication (Covaris S220 Focused Ultrasonicator) and library preparation with NEBNext Ultra II library preparation kit (NEB 7645).

Enrichment was carried out with a published set of biotinylated capture probes^14^. They were designed to be 120bp long with 50% overlap designed using the CapSequm tool (https://github.com/jbkerry/capsequm) (see Table S1 for details).

Hybridization and capture of targeted regions was carried out with the Roche HyperCapture Target Enrichment Kit (Roche 9075828001). First, 12ug of pooled ligated sequencing libraries and mouse COT DNA (10uL per individual library) were denatured by incubation at 95°C for 10min. Then the capture oligonucleotides were added (29 nM) and the hybridization reaction was carried out for 72h at 47°C. Hybridized DNA was isolated with M-270 Streptavidin Dynabeads, followed by washes and amplification. A second capture step was performed with a 24h hybridization time. Captured libraries were sequenced by 150 bp paired-end sequenced using a NovaSeq X Plus (Illumina).

### MCCu ANALYSIS

MCCu data was analysed with the published MCC pipeline^13^. Adapter sequences were removed with trim_galore (v0.3.1) and paired-end reads were merged using FLASH (v1.2.11). Initial alignment was carried out with the non-stringent aligner BLAT (v35) to identify the capture region (+800bp). The MCCuSplitter.pl script then divided the reads in the FASTQ file into sub-reads depending on the mapping by BLAT to the capture region and identified the position of sub-reads within the original reconstructed read. MCCuSplitter.pl was used to identify the two ligated fragments (capture viewpoint and interacting region) that constituted the read, divide them into subreads and output them into different FASTQ files depending on the capture region they mapped to. Bowtie2 (v2.3.5) was used to map sub-reads the mm39 reference genome. MCCuAnalyser.pl was used to stringently remove PCR duplicates and to identify the position of ligation junctions between all sub-reads as well as the orientation of the sub-read relative to the ligation junction.

For statistical analysis of long-range interactions, interchromosomal interactions were excluded and counts were normalised with the total number of *cis*-interactions. Ligation junctions were counted across active promoter regions (ATAC-seq peaks that were within 1kb of the TSS(NCBI RefSeq)), enhancers (ATAC-seq peaks overlapping with H3K27ac peaks located more than 1kb away from any TSS(NCBI RefSeq)), and CTCF sites (CTCF ChIP-seq peaks not classified as enhancers). Statistical significance of changes in interaction frequencies was assessed using the Wilcoxon rank-sum test, with *p*-values adjusted for multiple testing using the Benjamini–Hochberg procedure. Changes were considered significant if the adjusted *p-*value (padj) was < 0.05 and the absolute log2 fold change was ≥ 0.5. Fisher’s exact test was used to test the association between increased promoter–promoter interactions and transcriptional downregulation.

Raw contact matrices from ligation junctions were generated with cooler (v0.10.3). The python module iced^76^ was used for normalisation of symmetrical matrices. Matrices extended beyond the capture region were normalised to the total number of unique ligation junctions. Sequence reconstruction contact maps were generated as previously described^14^ and normalised to the total number of unique ligation junctions. In short, DNA binding proteins prevent MNase from cutting DNA at the protein bound sequences. The directionality of the read was determined whether the ligation junction is upstream or downstream to the protected DNA sequence. By leveraging the directionality of the reads from ligation junction pairs and extending them by a 10 × 4 bp rectangle, exact binding positions and interactions between DNA-associated proteins were determined. In addition, the Log2FC matrices for MCCu data were determined from sequence reconstruction contact maps (1bp res). The Log2FC square matrix was partitioned into consecutive, non-overlapping groups of 10 adjacent genomic bins that were aggregated by block-averaging to facilitate visualisation and reduce noise.

### Hi-C ANALYSIS

The Hi-C data for wildtype E14 cells were downloaded from GSE96107^52^. TAD boundaries were adapted to mm39 with liftOver. FASTQ files were processed using the HiC-Pro pipeline (v3.1.0)^53^. Reads were aligned to mm39. Default parameters were used to remove duplicates, assign reads to restriction fragments, filter valid pairs. Raw matrices were generated at 20000bp resolution. Replicates were merged and ICE normalised with cooler (v0.10.3). Contact matrices were generated for visualisation with CoolBox (v0.3.9).

### BIOINFORMATIC ANALYSIS OF ATAC-seq, CUT&Tag, ChIP-seq, PolyA RNA-seq, AND TT-seq

SeqNado (V0.7.6)^21^ was employed to process FASTQ files for ATAC-seq, CUT&Tag, ChIP-seq, polyA RNA-seq, and TT-seq data as described below.

ATAC-seq FASTQ files were quality checked using FastQC (v0.12.1). Adapter sequences were removed and low-quality reads were trimmed using trim_galore (v0.6.10) with the following parameters: --2colour 20. Reads were aligned to the reference genome mm39 using bowtie2 (v2.5.1) with the following parameters: --very-sensitive. Aligned reads were sorted by coordinate using samtools sort (v1.17). PCR duplicate reads were removed using Picard MarkDuplicates (v3.0.0) with the following parameters: REMOVE_DUPLICATES=true. Alignments were offset by +4bp (positive strand) and -5bp (negative strand) using bamnado (v0.3.11) to account for the Tn5 transposase adaptor insertions. BigWig files were generated using deepTools bamCoverage (v3.5.1) with the following parameters: --binSize 1 --normaliseUsing RPKM. Peak calling was performed using lanceotron (v1.2.6) with the following parameters: -c 0.5.

CUT&Tag FASTQ files were quality checked using FastQC (v0.12.1). Adapter sequences were removed and low-quality reads were trimmed using trim_galore (v0.6.10) with the following parameters: --2colour 20. Reads were aligned to a combined reference genome containing both mm39 (reference) and dm6 (spike-in) sequence using bowtie2 (v2.5.1) with the following parameters: --very-sensitive. Aligned reads were sorted by coordinate using samtools sort (v1.17). PCR duplicate reads were removed using Picard MarkDuplicates (v3.0.0) with the following parameters: REMOVE_DUPLICATES=true. Aligned reads were split into reference (mm39) and spike-in (dm6) fractions based on their alignment coordinates. Spike-in normalisation factors were calculated as previously described^77^ (1 / (spike-in reads / 1,000,000)). Data was analysed with both spike-in–based normalisation and conventional RPKM. While both approaches led to qualitatively similar conclusions, spike-in normalisation introduced increased variability across replicates of some samples (TBP, P300, CBP, RING1B, KDM2B, IgG). For these, we therefore report results based on RPKM normalisation, which provided more consistent signal across samples. BigWig files were generated using deepTools bamCoverage (v3.5.1) with the following parameters: --binSize 1 --normaliseUsing RPKM.

ChIP-seq FASTQ files were quality checked using FastQC (v0.12.1). Adapter sequences were removed and low-quality reads were trimmed using trim_galore (v0.6.10) with the following parameters: --2colour 20. Reads were aligned to a combined reference genome containing both mm39 (reference) and dm6 (spike-in) sequences using bowtie2 (v2.5.1) with the following parameters: --very-sensitive. Aligned reads were sorted by coordinate using samtools sort (v1.17). PCR duplicate reads were removed using Picard MarkDuplicates (v3.0.0) with the following parameters: REMOVE_DUPLICATES=true. Aligned reads were split into reference (mm39) and spike-in (dm6) fractions based on their alignment coordinates. Spike-in normalisation factors were calculated as previously described^77^ (1 / (spike-in reads / 1,000,000)). BigWig files were generated using deepTools bamCoverage (v3.5.1) with the spike-in derived normalisation factors and with --binSize 1. Peak calling was performed using lanceotron (v1.2.6) with the following parameters: -c 0.5.

For visualization of TT-seq, ATAC-seq, ChIP-seq, and CUT&Tag data, heatmaps and metaplots were generated with deepTools. Promoters were defined as region within 1kb a TSS (NCBI RefSeq) that overlap with ATAC-seq peak, and non-promoters as ATAC-seq peaks positioned >1kb from a TSS.

RNA-seq and TT-seq FASTQ files were quality checked using FastQC (v0.12.1). Adapter sequences were removed and low-quality reads were trimmed using trim_galore (v0.6.10) with the following parameters: --2colour 20. Reads were aligned to a combined reference genome containing both mm39 (reference) and dm6 (spike-in) sequences using STAR (v2.4.2a) with the following parameters: --quantMode TranscriptomeSAM GeneCounts --outSAMunmapped Within --outSAMattributes Standard. Aligned reads were sorted by coordinate using samtools sort (v1.17). Aligned reads were split into reference (mm39) and spike-in (dm6) fractions based on their alignment coordinates. Spike-in normalisation factors were calculated as previously described^77^ (1 / (spike-in reads / 1,000,000)). BigWig files were generated using deepTools bamCoverage (v3.5.1) with the spike-in derived normalisation factors and the following parameters: --binSize 1, --filterRNAstrand. For TT-seq, alignments were quantified over single merged transcripts (derived from NCBI RefSeq) using featureCounts (v2.0.6) with the following parameters: -t transcript -g gene_id -p --countReadPairs -s 2 -M -O. For PolyA RNA-seq, alignments were quantified over genes (NCBI RefSeq) using featureCounts (v2.0.6) with the following parameters -t exon -g gene_id -p --countReadPairs -s 2. Differential expression analysis was carried out with DESeq2 (v1.48.2) using the spike-in derived size factors and with the *padj* cutoff set to 0.05. For visualization MA plots were generated with ggplot2 and metaplots were generated with deepTools.

## SUPPLEMENTAL INFORMATION

**Document S1.** Figures S1-S4.

**Table S1**. Sequence and coordinates (mm39) of the capture oligonucleotides used for MCCu, related the STAR Methods.

