## Supplemental for "FACT safeguards promoter topology by maintaining nucleosomes and restricting chromatin factor spreading"

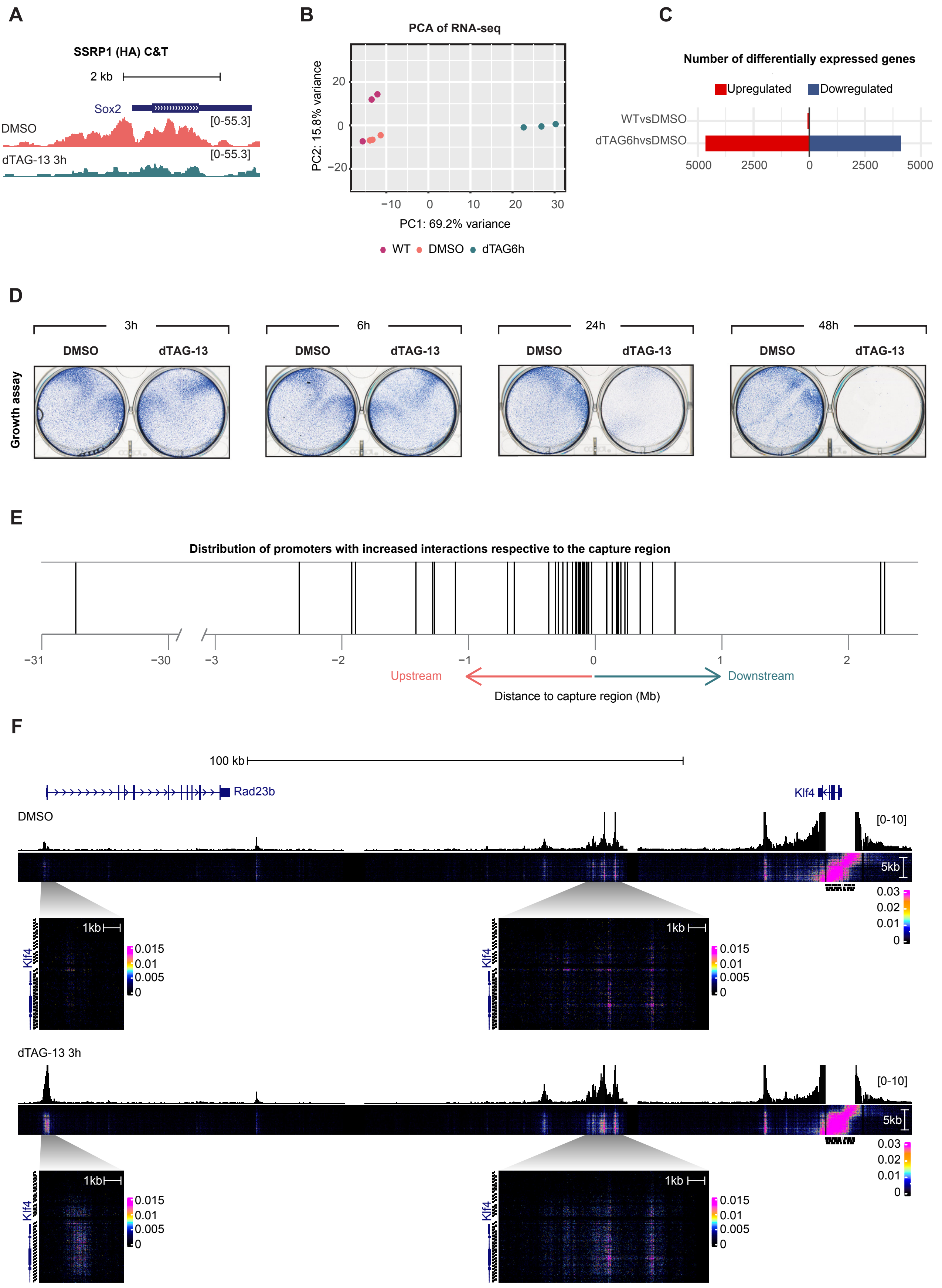

A
SSRP1 (HA) C&T
2 kb
Sox2
DMSO
[0-55.3]
dTAG-13 3h
[0-55.3]
B
PCA of RNA-seq
PC2: 15.8% variance
PC1: 69.2% variance
WT
DMSO
dTAG6h
C
Number of differentially expressed genes
Upregulated
Dowregulated
WTvsDMSO
dTAG6hvsDMSO
D
3h
6h
24h
48h
DMSO
dTAG-13
Growth assay
E
Distribution of promoters with increased interactions respective to the capture region
Upstream
Downstream
Distance to capture region (Mb)
F
100 kb
Rad23b
Klf4
DMSO
[0-10]
5kb
1kb
dTAG-13 3h
[0-10]

**Figure S1: Validation of the SSRP1-FKBP E14 cell line and further characterisation of increased promoter–promoter interactions, related to Figure 1.**

(A) CUT&Tag signal of SSRP1-FKBP (HA) at the *Sox2* locus in SSRP1-FKBP E14 cells treated with DMSO, and dTAG-13 for 3h. (B) PCA analysis of RNA-seq data shows that WT E14 and SSRP1-FKBP E14 cells treated with DMSO for 6h cluster together. By contrast, they clearly separate from SSRP1-FKBP E14 cells treated with dTAG-13 for 6h. (C) Barplot showing the number of differentially expressed genes between SSRP1-FKBP E14 cells treated with DMSO and, respectively, dTAG-13h 6h or WT cells. (D) Trypan blue growth assay of SSRP1-FKBP E14 cells treated with DMSO and 1μM dTAG-13 for 3h, 6h, 24h, and 48h. A representative replicate of three biological replicates is shown. (E) Distribution of distances between promoters with increased interactions and the corresponding capture promoter. (F) Extended MCCu heatmap outside the capture region (50bp resolution) and MCCu track (CPT normalised to *cis*-interactions) show an increase in interaction frequency and an expansion of the interacting surfaces upon FACT loss (chr4:55343700–55548600). Zoom-ins at regions of interest (chr4:55348000–55353000; chr4:55471480–55484020) are shown below (20bp resolution). The signal in contact matrices is normalised to the total number of unique ligation junctions and $\log_{10}$-transformed.

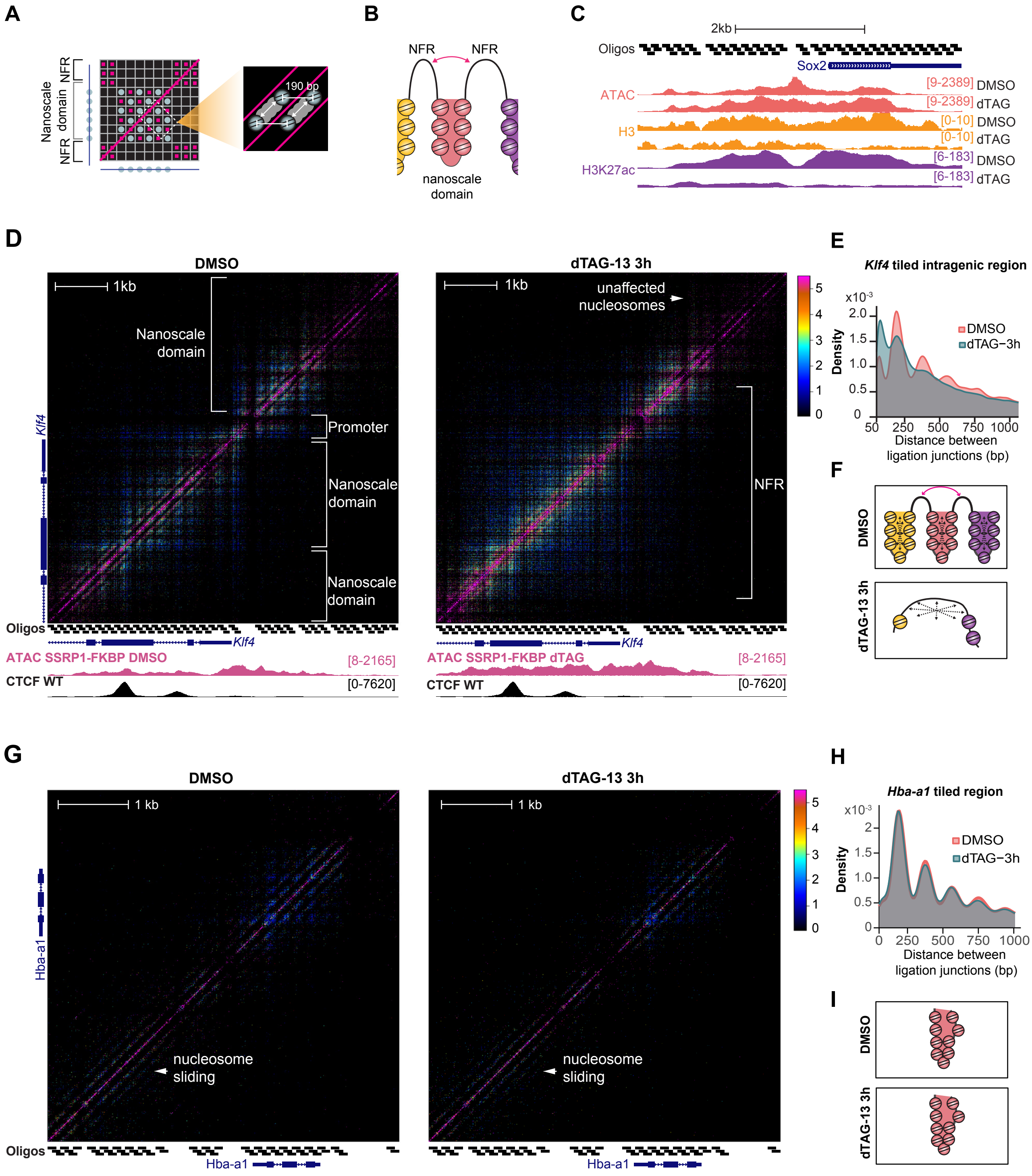

A
NFR
Nanoscale domain
NFR
190 bp
B
NFR
NFR
nanoscale domain
C
2kb
Oligos
Sox2
ATAC
[9-2389] DMSO
[9-2389] dTAG
H3
[0-10] DMSO
[0-10] dTAG
H3K27ac
[6-183] DMSO
[6-183] dTAG
D
DMSO
dTAG-13 3h
1kb
Nanoscale domain
Promoter
Nanoscale domain
Nanoscale domain
unaffected nucleosomes
NFR
Klf4
Oligos
ATAC SSRP1-FKBP DMSO
[8-2165]
CTCF WT
[0-7620]
ATAC SSRP1-FKBP dTAG
E
Klf4 tiled intragenic region
Density
DMSO
dTAG−3h
Distance between ligation junctions (bp)
F
DMSO
dTAG-13 3h
G
DMSO
dTAG-13 3h
1 kb
Hba-a1
nucleosome sliding
Oligos
H
Hba-a1 tiled region
Density
DMSO
dTAG−3h
Distance between ligation junctions (bp)
I
DMSO
dTAG-13 3h

**Figure S2: FACT depletion leads to an increase in interactions between distal promoters, related to Figure 2.**

(A) Schematic representation of an MCCu contact matrix. 3C contact signal is denoted by pink colouring. Nucleosomes positioning generates a stripping pattern where signal oscillates with a periodicity of ~190bp. (B) Model corresponding to the schematic MCCu matrix showing interactions within nanoscale domains, which are partitioned by nucleosome-free regions (NFRs). NFRs coalesce above aggregated nucleosomes and interact with neighbouring NFRs (C) ATAC-seq (RPKM), H3 ChIP-seq (ref-CPM) and H3K27ac (ref-CPM) at the *Sox2* locus supporting a decrease in nucleosomes upon FACT loss. (D) Contact matrix (5bp resolution, 6 replicates) at the *Klf4* promoter after treatment with DMSO, and dTAG-13 for 3h in SSRP1-FKBP E14 cells. Note the unperturbed nucleosomes upstream of the *Klf4* promoter. ICE was used to normalise the matrices of read junction density. Capture oligo positions, ATAC-seq (RPKM) for the DMSO control and dTAG-13 conditions are shown below the contact maps. CTCF ChIP-seq (CPM) from WT E14 is also shown. (E) Distribution of distances between ligation junctions in the intragenic tiled region of the *Klf4* locus comparing DMSO (red) and 3h dTAG-13 (teal). (F) Model of nano-scale domains in the *Klf4* locus, showing nucleosome depletion and loss of nanoscale domains upon FACT loss. (G) Contact matrix (5bp resolution) at the inactive *Hba-a1* promoter showing no noticeable changes after 3h of dTAG-13 treatment in SSRP1-FKBP E14. (H) Distribution of distances between ligation junctions in the inactive *Hba-a1* locus comparing DMSO (red) and 3h dTAG-13 (teal). (I) Model of nano-scale domains in the inactive *Hba-a1* locus, showing no changes in chromatin structure following FACT depletion.

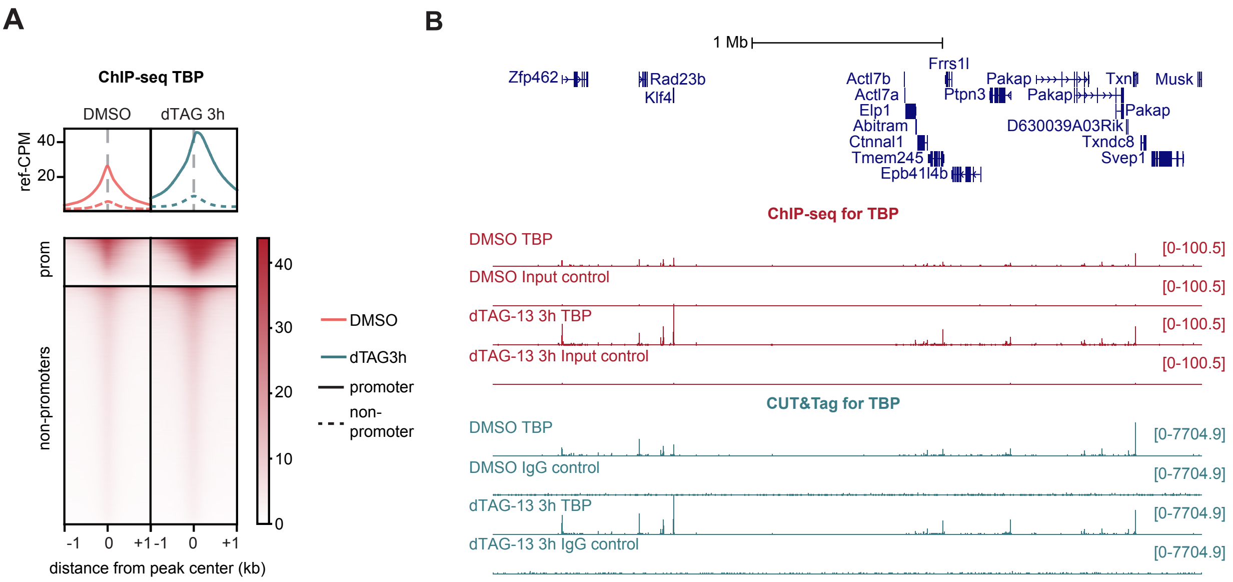
A
ChIP-seq TBP
DMSO
dTAG 3h
ref-CPM
40
20
prom
non-promoters
40
30
20
10
0
DMSO
dTAG3h
promoter
non-promoter
-1
0
+1
-1
0
+1
distance from peak center (kb)
B
1 Mb
Zfp462
Rad23b
Klf4
Actl7b
Actl7a
Elp1
Abitram
Ctnnal1
Tmem245
Epb41l4b
Frrs1l
Ptpn3
Pakap
Pakap
Pakap
D630039A03Rik
Txn
Txndc8
Svep1
Musk
ChIP-seq for TBP
DMSO TBP
[0-100.5]
DMSO Input control
[0-100.5]
dTAG-13 3h TBP
[0-100.5]
dTAG-13 3h Input control
[0-100.5]
CUT&Tag for TBP
DMSO TBP
[0-7704.9]
DMSO IgG control
[0-7704.9]
dTAG-13 3h TBP
[0-7704.9]
dTAG-13 3h IgG control
[0-7704.9]

**Figure S3: Orthogonal validation of TBP spreading, related to Figure 3.**

(A) Mean distribution and heatmap representation of ChIP-seq signal (ref-CPM) for TBP at promoters and other accessible regions (non-promoters) after treatment with DMSO (red) and 1μM dTAG-13 (teal). Promoters were defined as regions within 1kb a TSS (NCBI RefSeq) that overlap with ATAC-seq peak, and non-promoters as ATAC-seq peaks positioned >1kb from a TSS. (B) ChIP-seq (ref-CPM) and CUT&Tag (RPKM) signal for TBP and non-primary antibody controls around the *Klf4* locus.

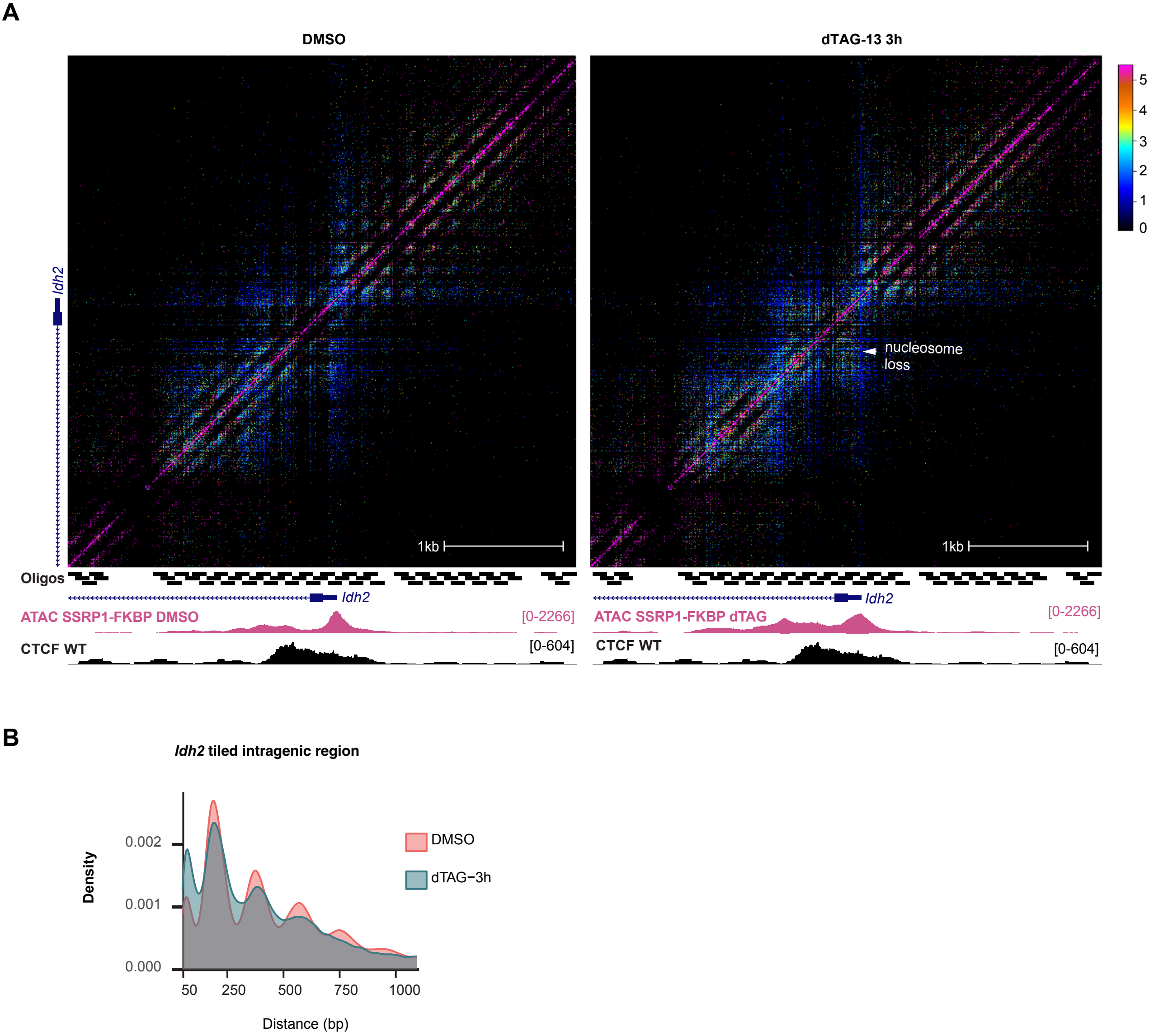

A
DMSO
dTAG-13 3h
Idh2
nucleosome loss
1kb
5
4
3
2
1
0
Oligos
Idh2
ATAC SSRP1-FKBP DMSO
[0-2266]
ATAC SSRP1-FKBP dTAG
CTCF WT
[0-604]
B
Idh2 tiled intragenic region
Density
0.002
0.001
0.000
50
250
500
750
1000
Distance (bp)
DMSO
dTAG−3h

**Figure S4**: **Disruption in local chromatin organisation in the *Idh2* locus upon FACT loss, related to Figure 5**

(A) Contact matrix (5bp resolution) at the *Idh2* promoter after treatment with DMSO, and dTAG-13 for 3h in SSRP1-FKBP E14 cells (ICE was used to normalise the matrices of read junction density; 6 replicates). Capture oligo positions, ATAC-seq (RPKM) for the DMSO control and dTAG-13 conditions are shown below the contact maps. CTCF ChIP-seq (CPM) from WT E14 is also shown. (B) Distribution of distances between ligation junctions in the tiled intragenic region of *Idh2* comparing DMSO (red) and 3h dTAG-13 (teal).
